# Differential Roles of PFDN5 Isoforms in Head and Neck Squamous Cell Carcinoma: Insights from Proximity Interactome Mapping

**DOI:** 10.64898/2026.05.07.723477

**Authors:** Chesnel Franck, Cheron Angélique, Audic Yann, Alusse Adrien, Duot Matthieu, Com Emmanuelle, Lavigne Régis, Paillard Luc, Le Goff Xavier

**Author notes:** current address: IRIBHM ULB, 808 route de Lennik, 1070 Brussels, Belgium.

## Abstract

Head and neck squamous cell carcinoma (HNSCC) ranks as the seventh most common cancer, with increasing incidence and mortality rates and limited therapeutic progress. The heterohexameric prefoldin complex, a highly conserved co-chaperone assembly composed of six PFDN subunits, exhibits expression levels strongly correlated with cancer progression. Among these subunits, the *PFDN5* gene presents a paradoxical role in cancer biology, demonstrating both tumor-promoting and tumor-suppressive activities. Notably, the *PFDN5* gene generates two distinct protein isoforms through alternative splicing, yet their individual contributions to cancer remain unexplored. In this study, we reveal that an elevated short-to-long *PFDN5* alternative splice variants ratio is significantly associated with improved overall survival in HNSCC patients. Using proximity-dependent biotin identification (BioID), we mapped shared and isoform-specific protein–protein interaction networks for PFDN5. Our analysis uncovered novel proximal interactors, implicating PFDN5 isoforms in unexpected functions, including spindle organization, transcriptional complexes, and NF-κB signaling. These results provide a foundation for exploring PFDN5 isoforms as potential therapeutic targets in HNSCC.

## INTRODUCTION

Prefoldin (PFD) is a highly conserved heterohexameric protein complex found in archaea and eukaryotes. This approximately 100 kDa co-chaperone was first identified biochemically in archaea and mammals, and genetically in yeast as the GIM complex (GIMc) (Geissler et al., 1998; Vainberg et al., 1998). In eukaryotes, PFD comprises two α-subunits (PFDN3 and PFDN5) and four β-subunits (PFDN1, PFDN2, PFDN4, and PFDN6), all of which share a conserved structural organization: NH_2_-terminal and COOH-terminal α-helical coiled-coil domains separated by either two (α-subunits) or one (β-subunits) β-hairpin motifs. When assembled, PFD adopts a jellyfish-like architecture (Vainberg et al., 1998). PFD plays a critical role in protein folding by delivering unfolded substrates, such as actin and tubulin, to the Cytosolic Chaperonin containing TCP-1 (CCT) complex (also known as the T-complex-protein-1-ring or TRiC) (Geissler et al., 1998; Gestaut et al., 2019; Hansen et al., 1999; Martín-Benito et al., 2002; Vainberg et al., 1998). PFD has been linked to cell migration through its regulation of β-actin folding (Fan et al., 2020). PFD has also been implicated in the folding of non-cytoskeletal substrates, including the human pVHL tumor suppressor and histone deacetylase 1 (HDAC1) (Banks et al., 2018; Chesnel et al., 2020). Additionally, PFD regulates the aggregation and degradation of misfolded proteins in yeast (Comyn et al., 2016) and has been shown to modulate protein aggregation in models of neurodegenerative diseases (Tashiro et al., 2013).

Beyond the canonical co-chaperone function of the PFD complex, individual PFDN subunits exhibit other PFD-independent roles (reviewed in (Herranz-Montoya et al., 2021). For example, PFDN1 promotes epithelial-mesenchymal transition (EMT) and cell invasion by regulating cyclin A transcription in the nucleus (Wang et al., 2017). PFDN5, also known as Myc Modulator-1 (MM-1), has been extensively studied for its tumor suppressor activity, primarily through inhibition of the c-MYC oncogene *via* multiple mechanisms (Fujioka et al., 2001; Hagio et al., 2006; Han et al., 2016; Kimura et al., 2007; Mori et al., 1998; Narita et al., 2012; Satou et al., 2004, 2001; Yoshida et al., 2008). However, the precise role of PFDN5 in cancer remains controversial, as evidence supports both tumor-promoting (Alldinger et al., 2005; Shao et al., 2024; Yesseyeva et al., 2020) and tumor-suppressing functions (Hennecke et al., 2015; Yu et al., 2024). Notably, PFDN5 may exert a dual role in breast cancer, inhibiting proliferation while simultaneously promoting cell migration and invasion (Wen et al., 2024). Nuclear PFDN5 has been implicated in RNA polymerase elongation, pre-mRNA splicing, and chromatin remodeling, functions that may be shared by some, but not all, PFDN subunits (Blanco-Touriñán et al., 2021; Millán-Zambrano et al., 2013; Payán-Bravo et al., 2021; Pells et al., 2015). The *PFDN5* gene was initially reported to encode four putative splice variants (Hagio et al., 2006), although only two are currently annotated in the GenBank database. The full-length mRNA variant is formed of 6 exons and encodes a 154-amino-acid protein, hereafter referred to as PFDN5α. A second splice variant, resulting from the skipping of exons 2 and 3, produces a 109-amino-acid isoform designated PFDN5γ (Fig. 1A).

**Figure 1:**
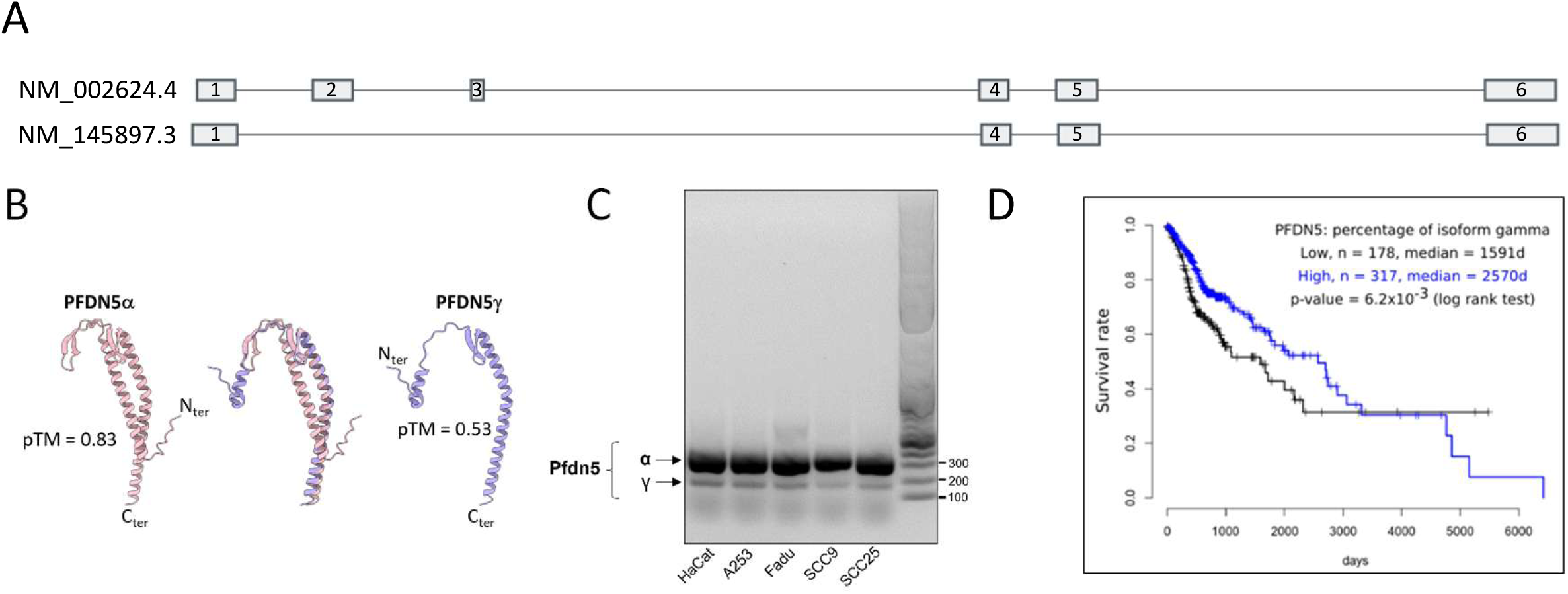
Expression and structural characterization of PFDN5 isoforms in head and neck squamous cell carcinoma (HNSCC). A) Gene organization of *PFDN5*: Two mRNA variants, NM_002624.4 (6 exons) and NM_145897.3 (4 exons), are generated from the *PFDN5* gene by alternative splicing. B) AlphaFold-predicted protein structures of PFDN5α and PFDN5γ isoforms, shown individually and in superposition. Per-chain confidence scores (pTM) are indicated. C) RT-PCR amplification of *PFDN5α* and *PFDN5γ* mRNA variants from total RNAs of HaCaT and four HNSCC cell lines (A253, FaDu, SCC9, SCC25). D) Kaplan-Meier survival analysis of 495 HNSCC patients stratified by the percentage of *PFDN5γ* mRNA (High: blue, *n*=317; Low: black, *n*=178), using data from the TCGA database.

Head and neck squamous cell carcinoma (HNSCC) ranks as the seventh most prevalent cancer worldwide, with approximately 800,000 new cases and 400,000 deaths annually (Sung et al., 2021). These tumors primarily originate from the epithelia of the oral cavity, nasal cavity, pharynx, larynx, salivary glands, and sinuses. HNSCC cancers are subdivided into two tumor subclasses: HPV-positive tumors and HPV-negative tumors. Tobacco smoking and alcohol consumption remain the predominant risk factors for HNSCC (Leemans et al., 2011). Notably, recent epidemiological data from the USA indicate a rising incidence and mortality for both oral cavity and pharyngeal HNSCC subtypes, contrasting with the declining trends observed in most other cancer types (Sherman et al., 2025). Current therapeutic strategies for HNSCC in addition to surgery, including radiotherapy, chemotherapy, and immunotherapy, are frequently undermined by drug resistance and radioresistance, limiting their efficacy. Despite these interventions, patient survival rates have stagnated over time, underscoring the urgent need for early detection biomarkers, prognostic tools, and innovative therapies.

The precise role of PFDN5 in tumorigenesis remains poorly understood. Notably, no study to date has investigated the distinct contributions of PFDN5 isoforms, generated through alternative splicing, to cancer progression. Here, we show by TCGA database analysis that elevated short-to-long isoform ratio correlates with improved overall survival in HNSCC patients. To elucidate the isoform-specific functions of PFDN5 and their potential involvement in HNSCC tumorigenesis, we employed proximity-dependent biotin identification (BioID), a powerful method for mapping *in cellulo* protein–protein interaction networks (Roux et al., 2012). Our findings significantly expand the known PFDN5 interactome, far beyond the interactions currently documented in databases such as BioGrid (Oughtred et al., 2021). We identified and confirmed novel shared and isoform-specific proximal proteins, pointing to unexpected functions for PFDN5 isoforms in spindle organization, post-transcriptional gene regulation, and epigenetic histone modifications.

## RESULTS

### 1) Expression levels of *PFDN5* alternative splice variants correlate with distinct outcomes in head and neck squamous cell carcinoma

The human *PFDN5* gene produces two mRNA variants by alternative splicing: *PFDN5α* (NM_002624.4) and *PFDN5γ* (NM_145897.3) (Figure 1A). *PFDN5α*, the full-length variant, encodes a 154-amino-acid protein with conserved structural features of α-class PFDN subunits, including NH_2_-terminal and COOH-terminal α-helical coiled-coil domains connected by a linker containing two β-hairpins (Figure 1B). In contrast, *PFDN5γ*, generated by skipping exons 2 and 3, encodes a 109-amino-acid isoform lacking one β-hairpin and featuring a truncated N-terminal α-helical domain (Figure 1B).

#### a) Differential Expression of *PFDN5* Isoforms in HNSCC Cell Lines

We assessed the amount of *PFDN5* isoforms in HNSCC cell lines using RT-PCR and RNA-seq. While both isoforms were detected in all samples, *PFDN5γ* mRNA levels were consistently and largely lower than *PFDN5α* in all cell lines (Figure 1C). Exploitation of GTEX data confirmed that tissues pertinent to HNSCC (Esophagus-Mucosa, Esophagus-muscularis, Esophagus-Gastroesophageal junction and Minor salivary Gland), had between 9 to 14.8 times more *PFDN5α* than *PFDN5γ*. In long read sequencing data from HaCaT cells representative of a keratinocyte cell line, the *PFDN5α* isoform is 23 times more represented than *PFDN5γ*. The tissue expressing the most *PFDN5γ* was the testis with only 2.5 times more *PFDN5α* than *PFDN5γ* isoform (supplementary Table S1).

#### b) The relative abundance of *PFDN5* isoforms has prognostic significance in HNSCC

We further examined the relative abundance of *PFDN5α* and *PFDN5γ* in HNSCC TCGA patients’ database. Strikingly, a high percentage of *PFDN5γ* correlates with improved overall survival (Fig. 1E). This does not rely on high expression of *PFDN5* gene, as global *PFDN5* RNA levels do not correlate with survival, confirming a previous report (Shao et al., 2024), and in contrast to other PFDN subunits (supplementary Figure S1).

### 2) Proximity-Dependent Interactome Analysis Reveals Common and Isoform-Specific PFDN5 Protein Networks

Cellular regulation and phenotypic diversity are ultimately determined by protein–protein interactions. To elucidate the isoform-specific interactomes of PFDN5, we employed an unbiased, proximity-dependent biotinylation approach (BioID) (Roux et al., 2012). This method uses a modified bacterial biotin ligase (BirA) fused to the protein of interest, enabling *in vivo* biotinylation of proximal proteins in living cells. Biotinylated proteins are then captured by affinity pulldown and identified by mass spectrometry. We transiently expressed BirA-tagged PFDN5 isoforms (PFDN5α-BirA-HA and PFDN5γ-BirA-HA) in human HNSCC FaDu cell line and conducted three independent BioID pull-down experiments (Supplementary Table S2). As controls, we included untransfected FaDu cells (treated with transfection reagent only) and FaDu cells expressing the BirA-HA ligase alone. We first confirmed the expression of all constructs BirA-HA, PFDN5α-BirA-HA, and PFDN5γ-BirA-HA by western blot (Figure 2A). Following 24-hour biotin supplementation, cells were lysed, and biotinylated proteins were affinity-purified using streptavidin-coated beads. The purified proteins were analyzed by TIMS-TOF Pro liquid chromatography-tandem mass spectrometry (LC-MS/MS), with peptide identification performed using Proline software.

**Figure 2:**
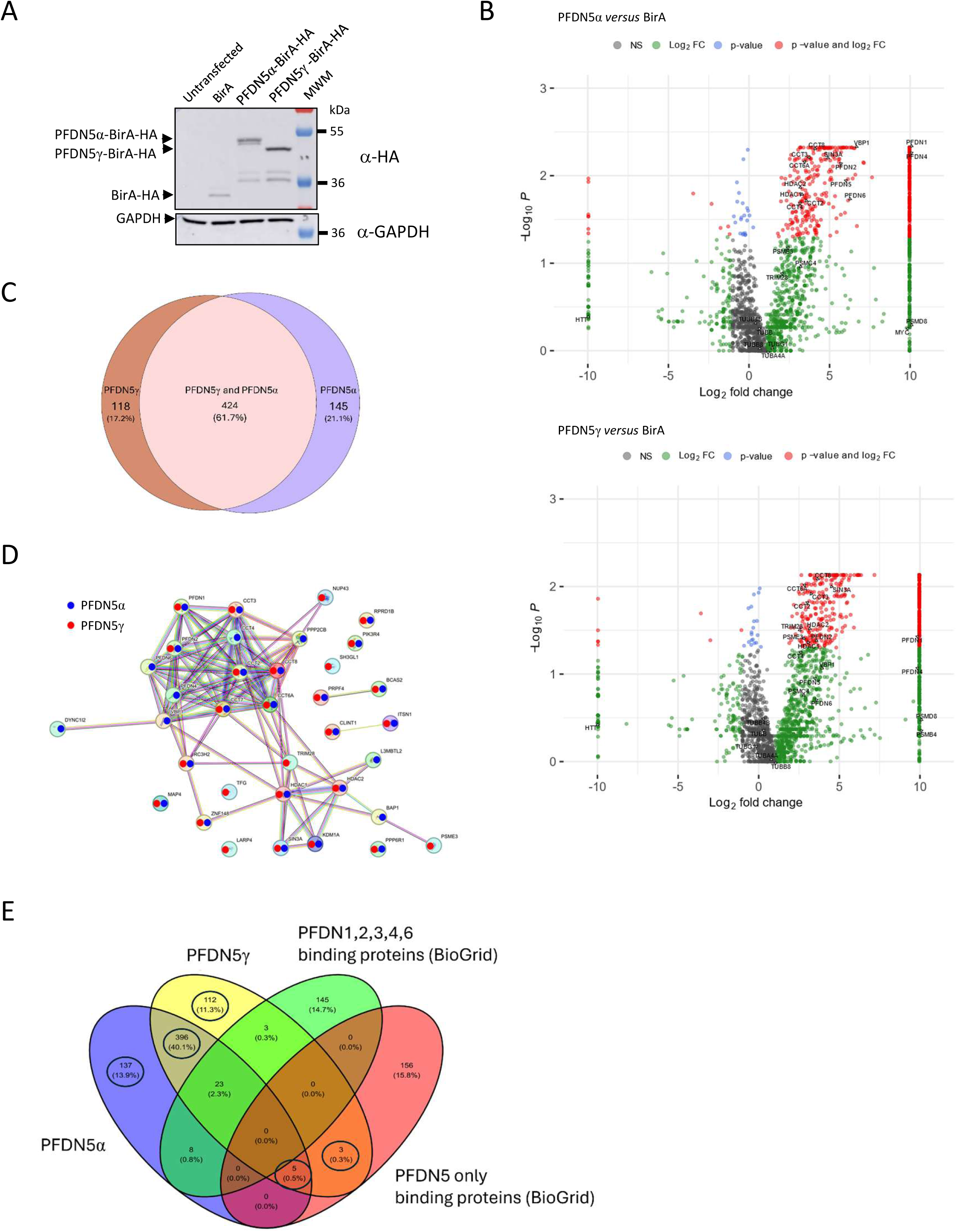
Isoform-specific PFDN5 interactomes identified by BioID. A) Immunoblot analysis of untransfected FaDu control cells or FaDu cells expressing HA-tagged BirA alone or fused to the indicated PFDN5 isoforms. GAPDH was used as a loading control. B) Volcano plots illustrating log₂ fold changes (LFC) in prey proteins between each PFDN5 isoform bait and control cells, as determined by BioID in FaDu cells. Significance was assessed using a beta-binomial test (5% false discovery rate, FDR; fold change threshold, *s₀*> 0.1). C) Venn diagram showing the overlap of interactome proteins identified by BioID for the indicated BirA-tagged PFDN5 isoforms. D) High confidence BioGrid PFDN5 interactors associated with PFDN5α, or PFDN5γ. E) Venn diagram showing the overlap of interactome proteins identified by BioID for the indicated BirA-tagged PFDN5 isoforms and PFDN interacting proteins from BioGrid.

### 2a) Statistical Analysis Reveals Isoform-Specific and Common PFDN5 Proximal Proteins

From the dataset of 5,492 identified proteins in any of the conditions, we applied a beta-binomial test to identify proteins with significantly different abundance in PFDN5α-BirA versus BirA (603 differential proteins) and PFDN5γ-BirA versus BirA (569 differential proteins), using multiple testing adjusted p-values (p < 0.05, Benjamini-Hochberg). To focus on biologically relevant interactions, we further filtered for proteins exhibiting a LFC > 1, yielding respectively 569 and 542 proteins with a higher recovery in PFDN5α-BirA or PFDN5γ-BirA compared to BirA only (Figure 2B and Supplementary Table S3).

Notably, these proteins enriched in PFDN5α or PFDN5γ samples included well-established PFDN5 partners, such as other PFDN subunits, CCT proteins, HDAC1, HDAC2 and SIN3A (Banks et al., 2018; Geissler et al., 1998; Satou et al., 2001; Vainberg et al., 1998), thereby providing a first line validation of the capacity of our BioID approach to capture *bona fide* PFDN5 interacting proteins. Confronting the PFDN5α-BirA and PFDN5γ-BirA enriched set (687 proteins total) revealed they shared 424 proteins (61.7% of the total), hereafter referred to as "PFDN5α-BirA and PFDN5γ-BirA common interactors" (Figure 2C), while 145 (21.1%) proteins were specific to PFDN5α and 118 (17.2%) to PFDN5γ (Figure 2C, Supplementary Table S4).

### 2b) Identification of characterized PFDN5 binding proteins

To compare with already known PFDN5α interacting proteins described in high throughput screen, we collected the interactome data for PFDNs proteins from the biomedical interaction repository BioGRid v4.4.241 (Oughtred et al., 2021). We selected only proteins demonstrating physical interactions with any of all six PFDN protein either as bait or as prey (Supplementary Figure S2, Table S5).

Analysis of the PFDN5α-BirA versus BirA and PFDN5γ-BirA versus BirA interactomes revealed that altogether 35 (5.1 % =35/687, (chi2(df=1,N=5371)=15.807, p= 7.014e-05 for the union of PFDN5α and PFDN5γ) proteins were present in the 146 BioGRid PFDN5-interacting proteins from the BioGrid database, 29 from PFDN5α-BirA versus BirA (4.8%, =29/569,(chi2(df=1,N=5371)=12.626, p= 0.0003804) and 27 from PFDN5γ-BirA versus BirA (4.7%, =27/542,(chi2(df=1,N=5371)=10.745, p= 0.001046). This indicated a significant enrichment of confirmed PFDN5 interacting protein in our BioID data both for PFDN5α and for PFDN5γ (Supplementary Tables S6,S7).

Interestingly, these 35 high confidence PFDN5 interactors are collectively mainly associated to the other prefoldin subunits, the CCT complex, and the SIN3A/HDAC1/HDAC2 complex. We can however detect some differences. While PFDN5α interacts with all PFDN subunits (1,2,VBP1/PFDN3, 4 and 6) PFDN5γ is only associated to PFDN1 and PFDN2. Among CCT complex proteins, only CCT4 appears to be preferentially associated to PFDN5α. BAP1, DYNC1l2, L3MBTL2 are also preferentially associated to PFDN5α. On the contrary, TRIM28, TFG, LARP4, NUP43, PSME3 and SH3GL1 are preferentially associated to PFDN5γ (Figure 2D). The presence of *bona fide* PFDN5 interacting proteins in our proteomic data validated our method. In addition, we identified approximately 95% of novel putative PFDN5 interactors.

### 2c) Identification of Putative Prefoldin Complex Substrate Proteins

The prefoldin complex is known to bind unfolded or misfolded proteins and transfer them to the CCT/TRiC chaperonin complex for proper folding. To date, only a limited number of prefoldin substrates have been characterized, including cytoskeletal proteins such as tubulin and actin monomers (Geissler et al., 1998; Hansen et al., 1999; Vainberg et al., 1998), the pVHL tumor suppressor (Chesnel et al., 2020), and the histone deacetylases HDAC1 and HDAC2 (Banks et al., 2018). However, these targets likely represent only a fraction of the proteins dependent on the prefoldin complex for proper folding.

With the aim to focus on PFDN5 specific functions, we seek to distinguish prefoldin substrate proteins (PSPs) from specific PFDN5-interacting proteins. We reasoned that PSPs should be associated to multiple PFDN subunits while proteins associated only to PFDN5 could be involved in the non-canonical functions or isoform-specific functions of PFDN5.

We therefore used BioGRid data collected above to filter out proteins associated to any PFDN subunit other than PFDN5 (Figure 2E). 179 proteins were associated to any of PFDN1,2,3,4 or 6 in BioGRid data of which 3 were associated also to PFDN5γ, 23 to both PFDN5α and PFDN5γ and 8 to PFDN5α only. These 34 proteins most likely represent PSPs or component of the PFDN and CCT complex. Among these 34 proteins interacting with PFDN5 isoforms and other PFDN, CCT8, CCT3, CCT7, CCT2, CCT6A, CCT4 are component of the CCT complex; PFDN1, PFDN2, VBP1, PFDN4,PFDN5,PFDN6 are component of the prefoldin complex; RUVBL2, RUVBL1, PIH1D1, RPAP3 are components of the PAQosome (Lynham and Houry, 2018). Other proteins associated to PFDN1,2,3,4,6 in BioGrid data were HDAC1, PRPF4, RC3H2, ARGHAP32, BAP1, PPP2CB, PIK3R4, DYNC1L2 for PFDN5α-BirA and HDAC1, PRPF4, RC3H2, ARGHAP32, PIK3R4, NUP43, PPP6R1 for PFDN5γ-BirA.

### 2d) Identification of PFDN5-Specific Interacting Proteins

In the BioGRID database, 164 proteins were uniquely associated with PFDN5. Among those PFDN5 specific interactant in BioGRID, only 8 proteins were exclusively associated to PFDN5 in BioID. Among these, five proteins were detected as proximal interactors of both PFDN5α-BirA and PFDN5γ-BirA in the BioID analysis and were confirmed to interact only with PFDN5 in BioGRID: SIN3A, KDM1A, ITSN1, BCAS2, ZNF148. Additionally, three proteins were identified as PFDN5γ-BirA-specific proximal interactors in the BioID dataset and were also found to interact exclusively with PFDN5 in BioGRID: PSME3, TRIM28, TFG. These findings suggest that these proteins are potentially involved in PFDN5-specific functions.

In a broader view, a total of 137 proteins were only associated to PFDN5α, 115 (112+3) were associated to PFDN5γ only and 401 (396+5) were commonly associated to PFDN5 alpha and gamma but to no other prefoldin subunit (Figure 2E).

### 3) The PFDN5*α* Isoform, but Not PFDN5*γ*, incorporates into the PFD Complex to Fulfill Prefoldin-Dependent Folding Functions

#### 3a) Structural Prerequisites for PFD Complex Incorporation

The α-class PFDN subunits (PFDN3 and PFDN5) dimerize via their extra β-hairpins, forming the structural core required for PFD hexamer assembly. Although PFDN5γ appears to retain a β-hairpin, this protein does not exhibit the characteristic structure of β-class PFDN subunits (PFDN1, 2, 4 and 6) (Siegert et al., 2000). Thus, we hypothesized that truncation in the NH_2_-terminal region of PFDN5γ might prevent its incorporation into the PFD complex. To test this, we first used AlphaFold 3.0 to model the PFD complex containing either PFDN5α or PFDN5γ alongside the other five PFDN subunits. The interface predicted template modelling ipTM score measures the accuracy of the predicted relative positions of the subunits forming the protein-protein complex. The ipTM score was lower for PFDN5γ (ipTM = 0.64) compared to PFDN5α (ipTM = 0.72), suggesting that PFDN5γ fails to efficiently integrate into the PFD complex (Figure 3A). Structural inspection revealed that the missing β-hairpin in PFDN5γ hinders critical interactions with neighboring PFDN1 and PFDN3 subunits (Figure 3A, inserts).

**Figure 3:**
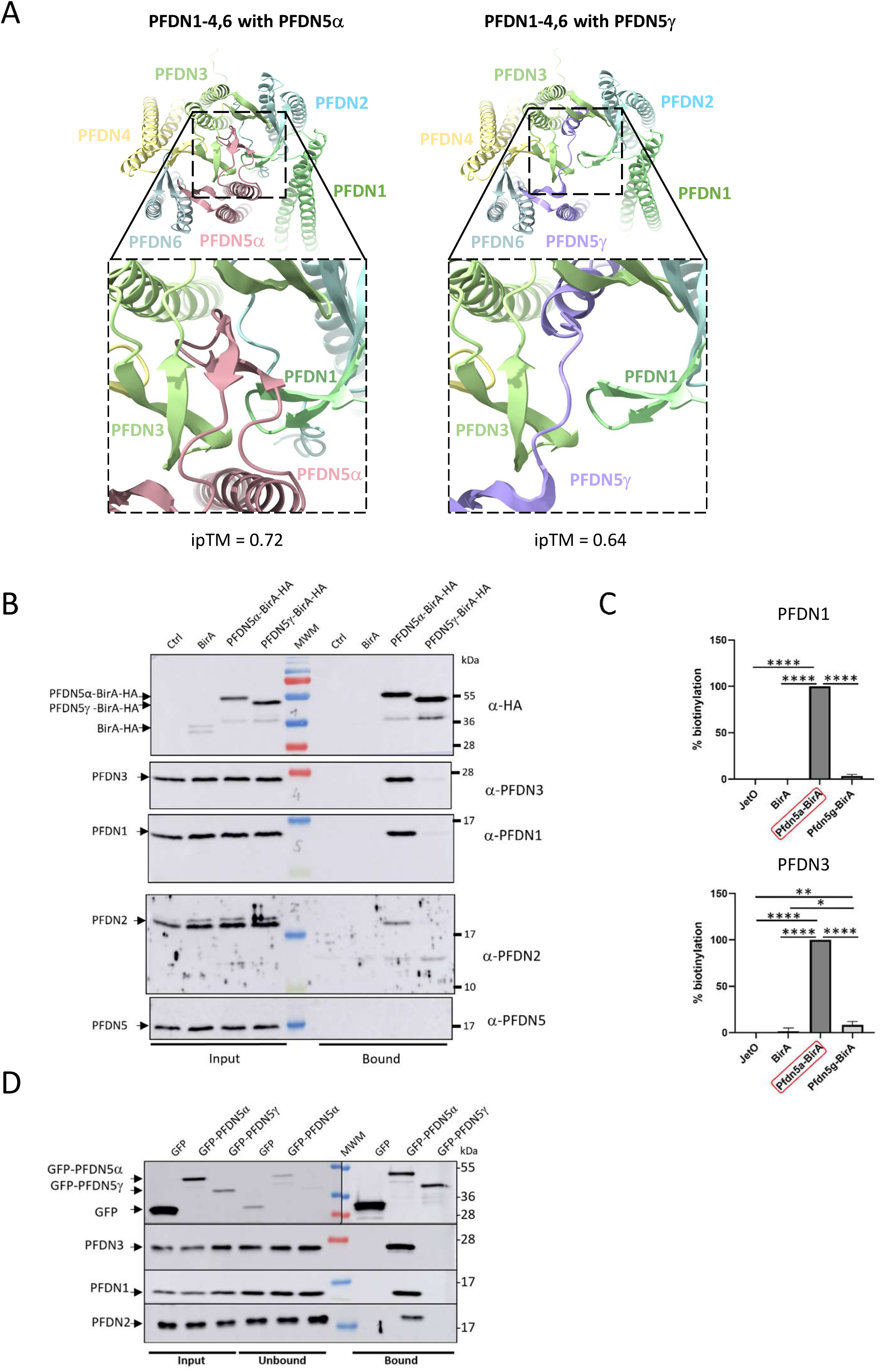
PFDN5α, but not PFDN5*γ*, associates with other prefoldin subunits. A) AlphaFold predictions of the interactions between each PFDN5 isoform and the five other PFDN (1-4,6) subunits. Interface predicted template modelling ipTM scores are indicated. B) Immunoblot analysis of HA-tagged BirA alone or fused to the indicated PFDN5 isoforms, or other PFDN subunits before (Input, left) and after (Bound, right) affinity capture using streptavidin-conjugated beads. Ctrl, untransfected control cells. C) Quantification of the biotinylated proteins (bound proteins shown in B) normalized against the level of expressed BirA alone or PFDN5s-BirA (Input proteins shown in B); * p<0.05, **, p<0.005, ****p<0.0001. D) Co- Immunoblotting of PFDN1, PFDN2, and PFDN3 after immunoprecipitation of GFP and GFP-tagged PFDN5 proteins using GFP-trap. MWM, Molecular weight pre-stained markers.

#### 3b) Experimental Validation of PFDN5 Isoform Incorporation

To experimentally validate these predictions, we performed BioID pull-down assays in FaDu HNSCC cells expressing PFDN5α-BirA or PFDN5γ-BirA fusion proteins. Western blot analysis of the biotinylated protein fractions showed that glyceraldehyde-3-phosphate dehydrogenase (GAPDH), a housekeeping enzyme localized in both the cytoplasm and nucleus, was not detected in the bound fractions of either PFDN5 isoform, serving as a negative control to validate the specificity of our results (see below, Figure 7B). Furthermore, the PFDN5α-BirA and PFDN5γ-BirA proteins were found in the bound fraction due to their auto-biotinylation. Strikingly, PFDN1, PFDN2, and PFDN3 co-eluted with PFDN5α-BirA, confirming their incorporation into the PFD complex (Figure 3B,C). In stark contrast, less than 10% of these PFDN subunits co-eluted with PFDN5γ-BirA (Figure 3B,C), indicating that PFDN5γ is largely excluded from the PFD complex. Notably, endogenous PFDN5 was absent from the PFDN5α-BirA pull-down, consistent with the stoichiometry of one PFDN5 subunit per PFD complex. These results were independently confirmed in A253 HNSCC cells (Supplementary Figure S3). To further corroborate these findings, we conducted co-immunoprecipitation experiments using GFP-tagged PFDN5 isoforms expressed in FaDu cells. PFDN1, PFDN2, and PFDN3 specifically co-immunoprecipitated with GFP-PFDN5α but not with GFP-PFDN5γ. No PFDN protein was found to be co-immunoprecipitated with GFP alone despite a much higher abundance than that of GFP-PFDN5 fusion proteins (Figure 3D). Collectively, these data demonstrate that PFDN5α, but not PFDN5γ, incorporates into the PFD complex, likely due to structural constraints imposed by the truncated NH_2_-terminal region of PFDN5γ.

#### 3c) Functional Rescue of PFD-Dependent Folding in Fission Yeast

Alpha- and beta-tubulins are conserved substrates of the PFD-CCT folding pathway, and PFD inhibition disrupts microtubule organization, leading to phenotypes such as hypersensitivity to thiabendazole (TBZ) and cold sensitivity in fission yeast (Chesnel et al., 2020). The fission yeast *pfd5* gene is the ortholog of human *PFDN5*, sharing 45.2% sequence identity with PFDN5α and retaining all structural features of an α-class PFDN subunit (Chesnel et al., 2020).

To assess whether PFDN5γ could functionally substitute for PFDN5α in the PFD complex, we expressed human PFDN5α or PFDN5γ in a fission yeast *pfd5*Δ deletion mutant using a constitutive pADH415 plasmid. While PFDN5α or GFP-PFDN5α (or PFDN5α-BirA, data not shown) expression rescued the growth defect of *pfd5*Δ cells on TBZ-containing plates and at 20°C, PFDN5γ expression failed to restore growth, phenocopying the empty vector control (Figure 4A).

**Figure 4:**
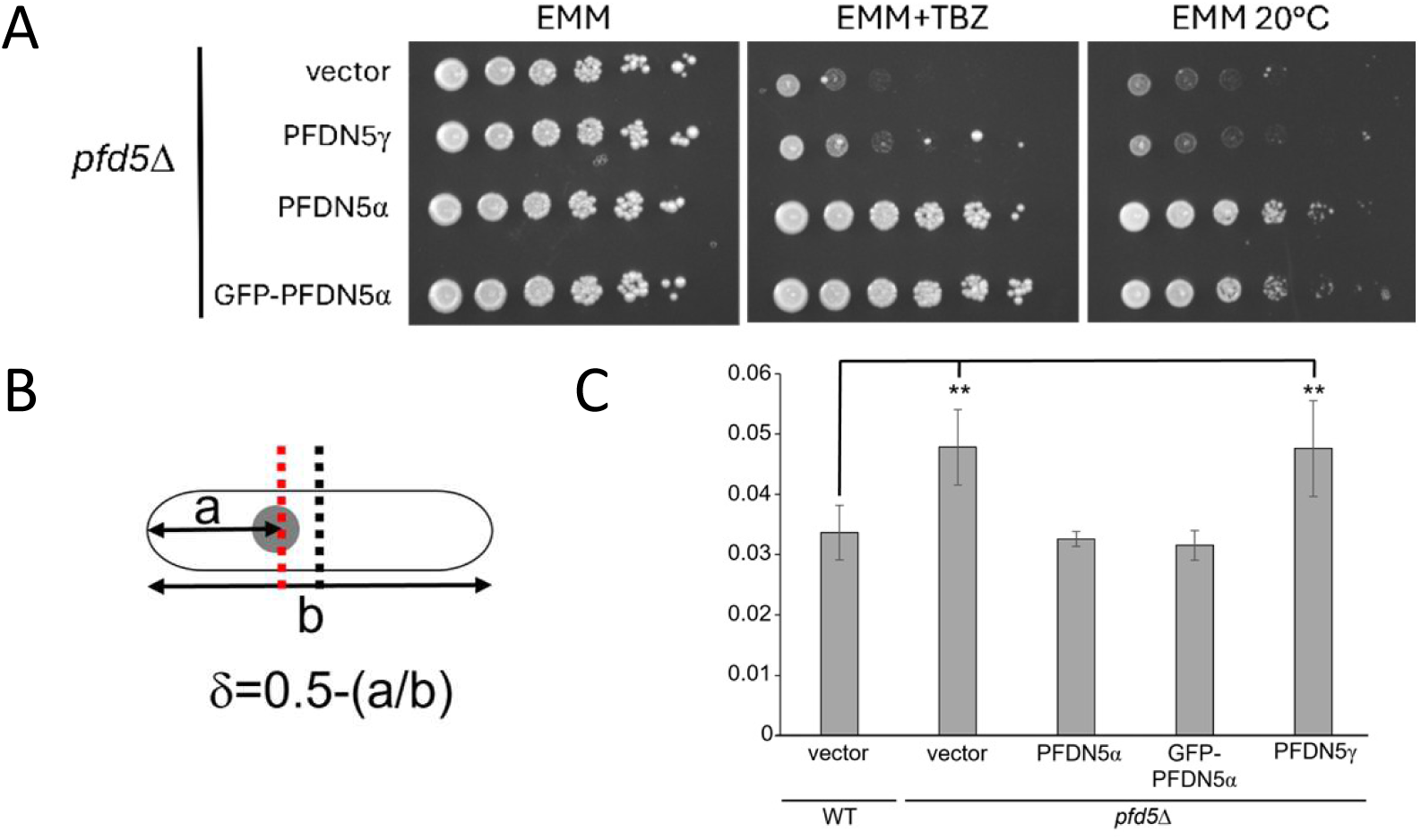
**PFDN5α, but not PFDN5*γ*, rescues *pfd5***Δ **fission yeast mutant phenotype.** A) Growth assay of fission yeast strains transformed with plasmids expressing the indicated proteins (empty vector as negative control). Serial dilutions (1:5) were spotted on EMM (control) or EMM+TBZ plates and incubated for 3 days at 25°C or 5 days at 20°C. B) Schematic representation of nuclear positioning (a) analysis in fission yeast cells, relative to the geometric cell center (b). C) δ values are reported in the histogram for wild-type (WT) and *pfd5*Δ strains (*n* = 30 cells per condition; mean δ values from 3 independent experiments; *p* < 0.05 (*) and *p* < 0.01 (**)).

In wild-type fission yeast, the interphase nucleus is maintained at the geometric cell center by balanced microtubule-dependent pushing forces. PFD-deficient cells (*pfd5*Δ) exhibit asymmetric nuclear positioning due to defective tubulin folding (Chesnel et al., 2020). To quantify this phenotype, we measured the nucleus-to-cell-end distance ratio in interphase cells and calculated δ, the deviation from the theoretical centered position where δ = 0 (Figure 4B). In wild-type cells, the mean δ was 0.033 ± 0.004. In contrast, *pfd5*Δ cells exhibited a significantly increased δ (0.047 ± 0.002, p = 3.6x10^-3^), reflecting nuclear mispositioning. Strikingly, expression of human PFDN5α fully rescued this phenotype (δ = 0.032 ± 0.007), as did the fusion protein GFP-PFDN5α (δ = 0.031 ± 0.005). In contrast, PFDN5γ expression failed to restore nuclear centering (δ = 0.047 ± 0.006, p = 0.04189), mirroring the *pfd5*Δ mutant phenotype (Figure 4C, Supplementary Figure S4). These results confirm that PFDN5α, but not PFDN5γ, rescues the PFD-dependent folding function in fission yeast, likely by restoring proper tubulin folding and microtubule dynamics.

### 4) PFDN5*α* and PFDN5*γ* Isoforms Exhibit Dual Cytoplasmic and Nuclear Localization

#### 4a) Rationale for Investigating Nuclear Localization

While the canonical function of the PFD complex in neosynthesized protein folding is thought to occur primarily in the cytoplasm, emerging evidence suggests that PFDN subunits, including PFDN5, may also localize to the nucleus. Previous studies in yeast, plant, and human cells have reported nuclear functions for PFDN5 (Blanco-Touriñán et al., 2021; Millán-Zambrano et al., 2013; Payán-Bravo et al., 2021; Pells et al., 2015). Our proximity interactome analysis further revealed interactions with nuclear proteins, prompting us to investigate the subcellular localization of PFDN5 isoforms in greater detail.

#### 4b) Dual Localization of PFDN5 Isoforms in FaDu Cells

To examine the localization of PFDN5α and PFDN5γ, we expressed GFP-tagged isoforms in human FaDu cells. Fluorescence confocal microscopy revealed that both isoforms localized to the cytoplasm and the nucleus (Figure 5A). This dual localization was independently confirmed by indirect anti-HA immunofluorescence using PFDN5α-HA-transfected FaDu cells (Supplementary Figure S5).

**Figure 5:**
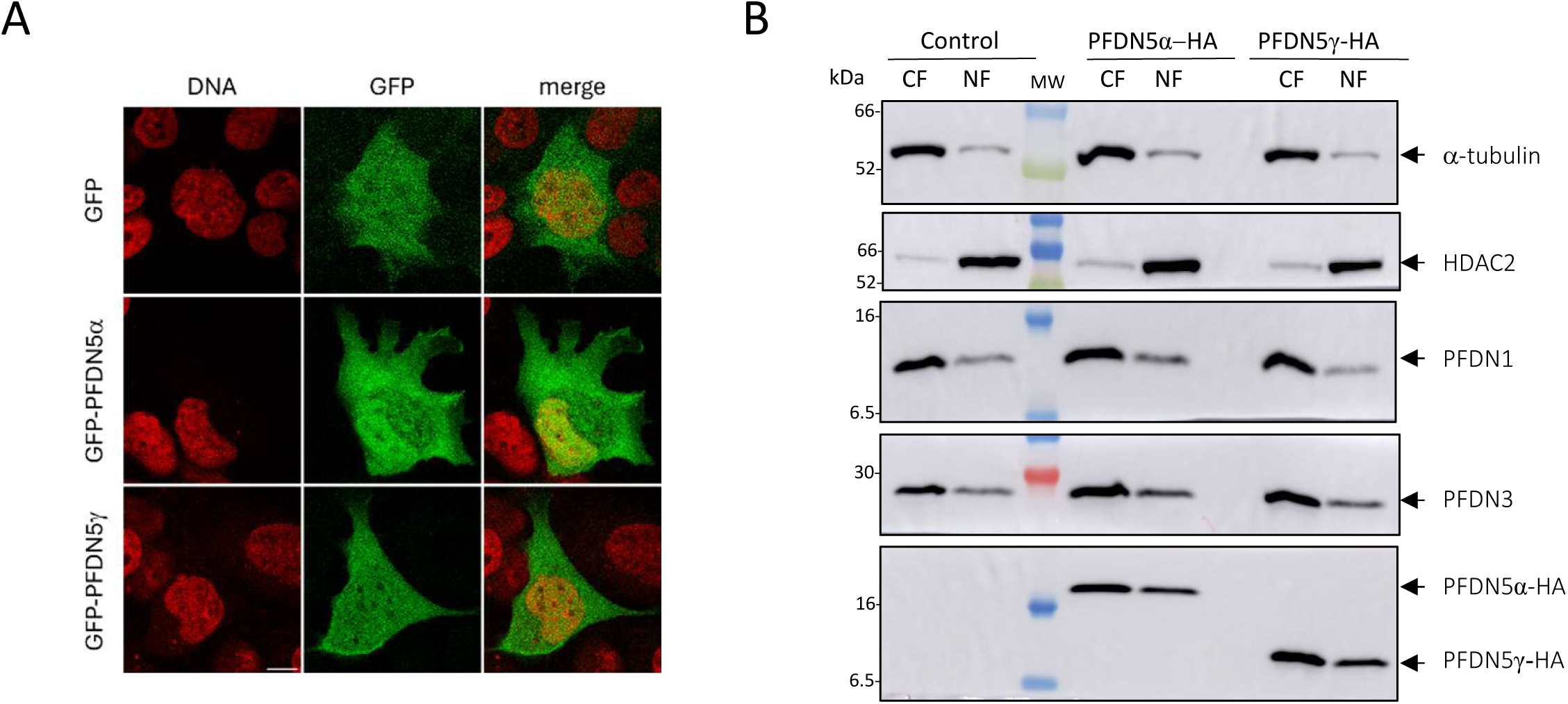
Nuclear localization of PFDN5 isoforms. A) Representative confocal microscopy images of FaDu cells showing the nuclear colocalization of GFP-tagged PFDN5 isoforms with DAPI labeled nuclei. GFP alone was used as a control. Scale bar, 10 µm. B) Western blot analysis of subcellular fractions from FaDu cells expressing PFDN5α-HA or PFDN5γ-HA. Protein extracts were separated into cytosolic (CF) and nuclear (NF) fractions. Target proteins are indicated on the right. MW, Molecular weight pre-stained markers.

#### 4c) Biochemical Fractionation Confirms Nuclear Localization of PFDN5 Isoforms

To biochemically validate these observations, we performed subcellular fractionation of FaDu cells, separating cytoplasmic (CF) and nuclear (NF) fractions. As expected, the cytoplasmic marker α-tubulin was largely enriched in the CF fraction while the nuclear marker HDAC2 was predominantly detected in the NF fraction (control, Figure 5B). Consistent with their cytoplasmic role in protein folding, endogenous PFDN1 and PFDN3 were primarily found in the CF fraction (control, Figure 5B; 73±10% and 76±6, respectively, Table 1). To specifically detect PFDN5 isoforms, we generated C-terminal HA-tagged constructs under the control of low-expression (pPGK) or high-expression (pCMV) promoters. We selected pPGK-PFDN5α-HA and pCMV-PFDN5γ-HA plasmids, as their transfection resulted in comparable expression levels of recombinant proteins (Supplementary Figure S6). Following fractionation of transfected FaDu cells, both PFDN5α-HA and PFDN5γ-HA were predominantly detected in the CF fraction: 73±7% and 71±15% for PFDN5α and PFDN5γ, respectively (middle and right panels, Figure 5B; Table 1). However, a significant proportion of each isoform was also present in the NF fraction (Figure 5B; 27±7% and 29±15% for PFDN5α and PFDN5γ, respectively, Table 1). These proportions were significantly higher than the expected leakage during the subcellular fractionation protocol, as inferred from the specific markers α-tubulin and HDAC2 used as controls in the experiment (<10%, Table 1). This result confirmed the dual subcellular localization of PFDN5α and PFDN5γ in the cytoplasm and in the nucleus.

**Table 1:**
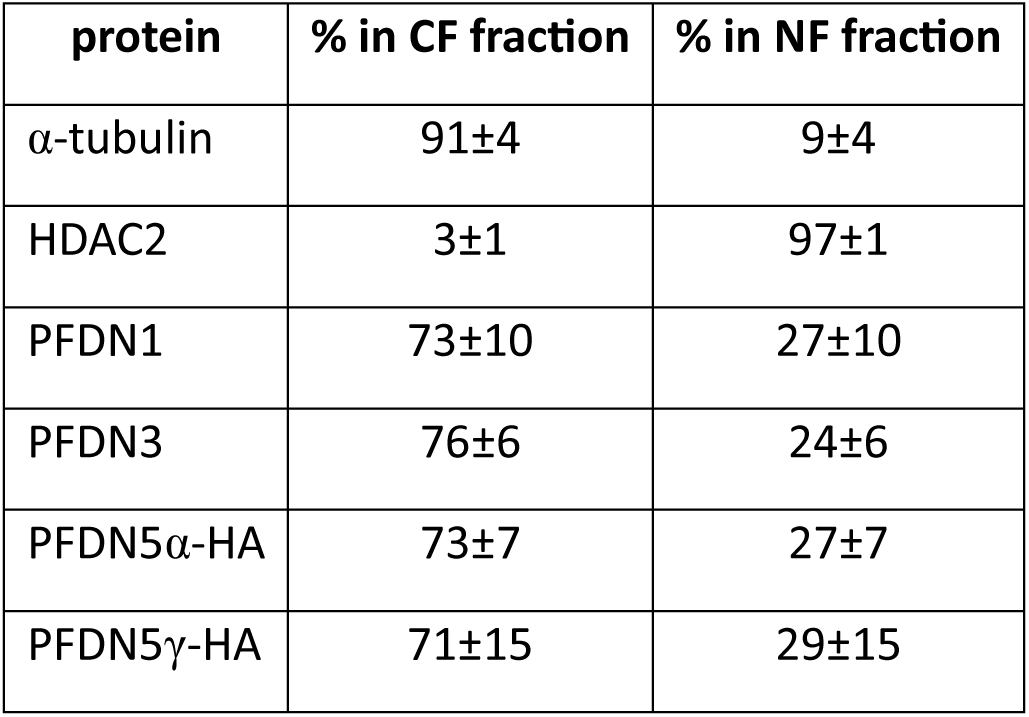
Quantification of protein distribution in subcellular compartments (n=4)

| **protein** | **% in CF fraction** | **% in NF fraction** |
| --- | --- | --- |
| α-tubulin | 91±4 | 9±4 |
| HDAC2 | 3±1 | 97±1 |
| PFDN1 | 73±10 | 27±10 |
| PFDN3 | 76±6 | 24±6 |
| PFDN5α-HA | 73±7 | 27±7 |
| PFDN5γ-HA | 71±15 | 29±15 |

### 5) Identification of Protein Complexes in the Proximity Interactomes of both PFDN5*α* and PFDN5*γ* by CORUM Enrichment

We reasoned that by the proximal ligation of biotin to complexes loosely associated to PFDN5α or PFDN5γ, our BioID experiment should label multiple components of any complexes in proximity to the PFDN5 isoforms. Therefore, we queried the CORUM database (Tsitsiridis et al., 2023)to identify complexes with overrepresented components in any subset of our BioID experiments.

Altogether, six main protein interaction networks (PINs) were enriched in the union of PFDN5α and PFDN5γ BioID targets (Figure 6A). They corresponded to expected complexes such as 1) the prefoldin complex or 2) the CCT/TRiC chaperonin complex but also to less expected ones such as 3) the HAUS augmin-like complex (Lawo et al., 2009; Uehara et al., 2009), to 4) multiple interconnected complexes where HDAC1 was central (Asmamaw et al., 2024), to 5) subunits of the 26S proteasome (Kanayama et al., 1992) and to the R2TP/prefoldin-like complex (Lynham and Houry, 2018).

**Figure 6:**
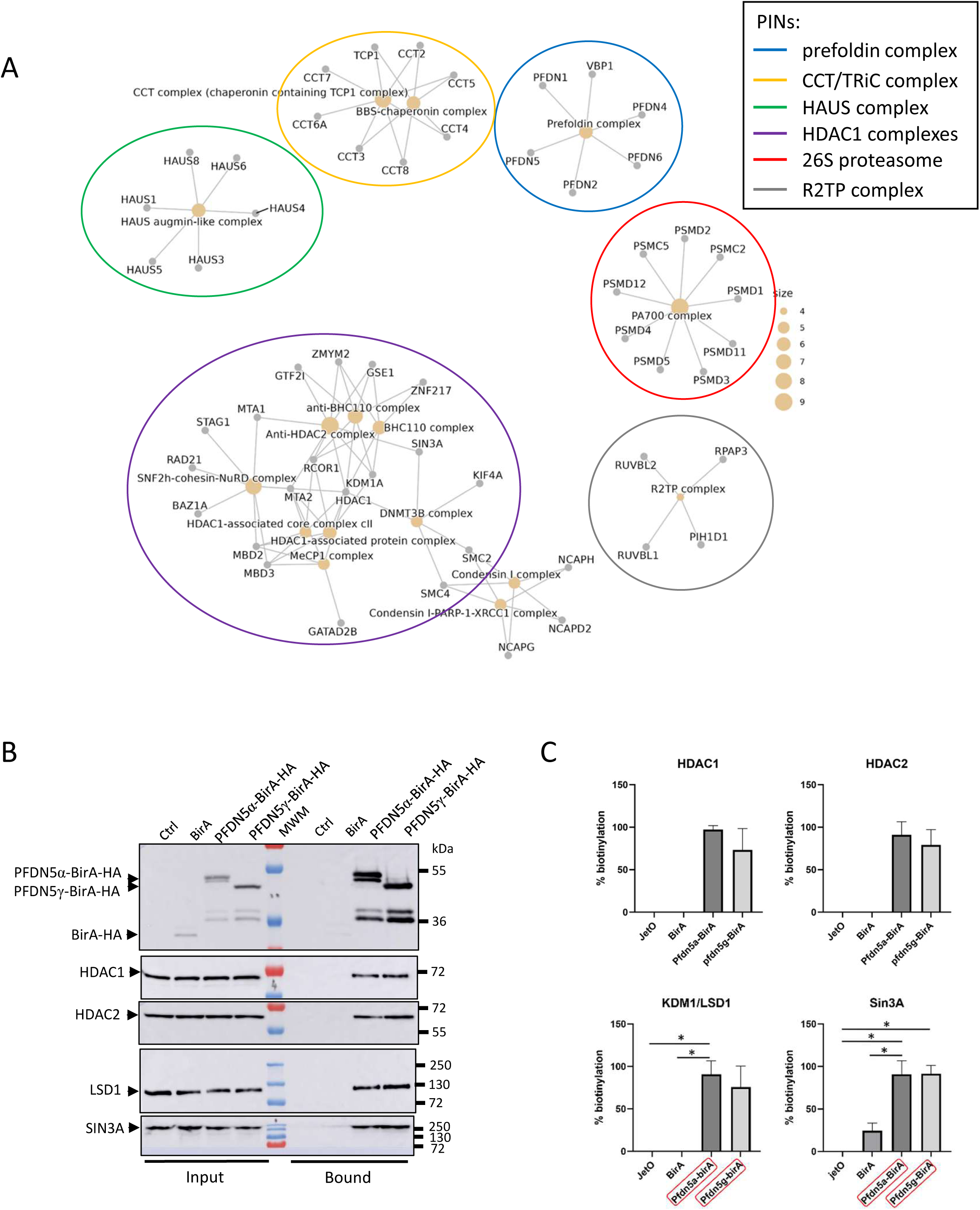
Identification of PFDN5 isoform-interacting proteins in FaDu cells using BioID and LC-MS/MS. A) Proteins identified in the most represented CORUM categories (alpha_or_gamma). Six main protein interaction networks (PINs) are indicated by a color code. (B) Immunoblot analysis of HA-tagged BirA alone or fused to the indicated PFDN5 isoforms, HDAC1, HDAC2, LSD1, and SIN3A, before (Input, left) and after (Bound, right) affinity capture using streptavidin-conjugated beads. Ctrl, untransfected control cells. MWM, Molecular weight pre-stained markers. C) Quantification of the biotinylated proteins (bound proteins shown in B) normalized against the level of expressed BirA alone or PFDN5s-BirA (Input proteins shown in B); * p<0.05.

**Figure 7:**
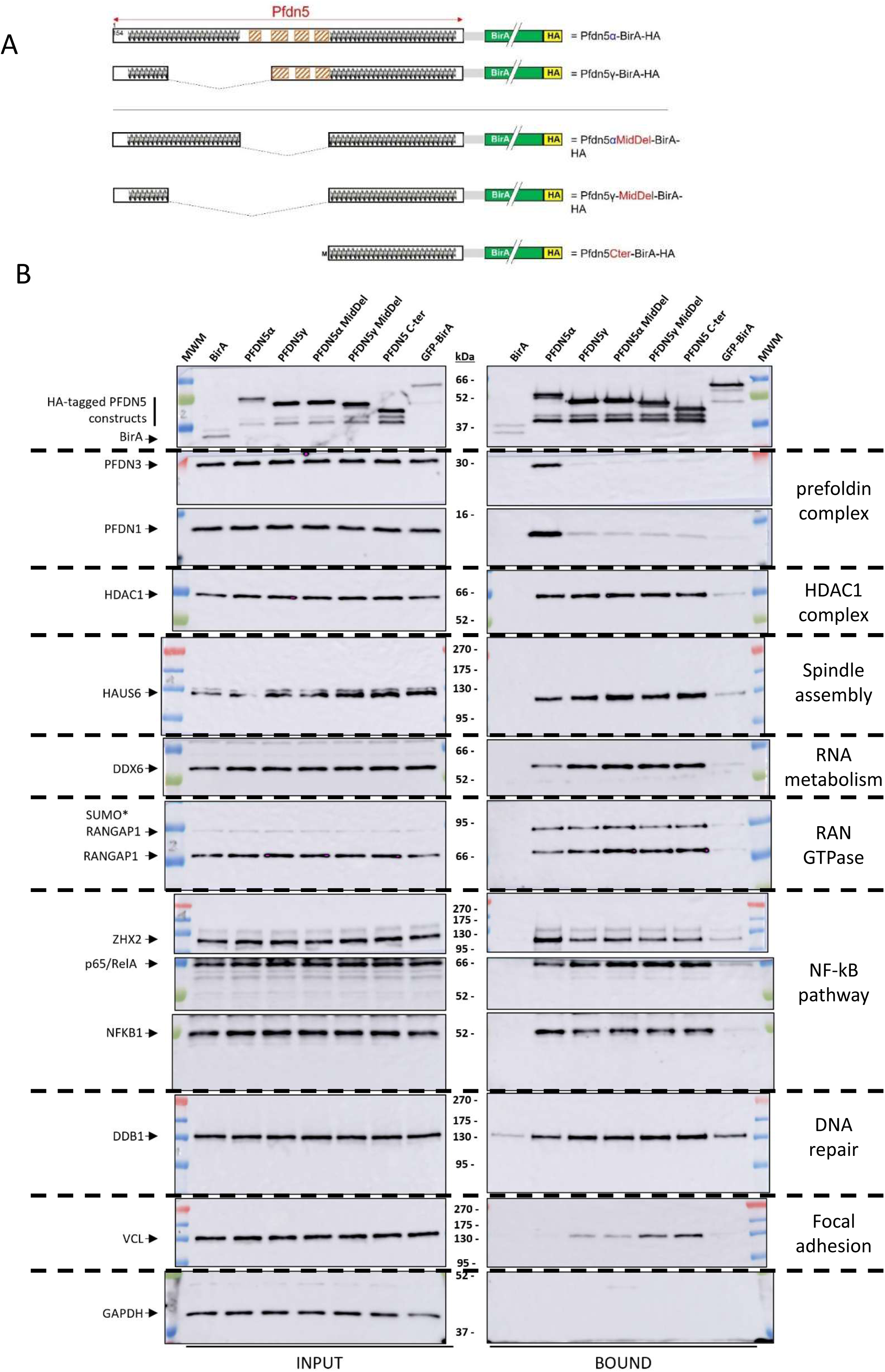
**Mapping the interactions of candidate protein partners with PFDN5 isoforms**. A) Schematic representation of BirA-HA-tagged truncated PFDN5 constructs. B) Immunoblot analysis of HA-tagged BirA alone or fused to the indicated PFDN5 isoforms and candidate target proteins, before (Input, left) and after (Bound, right) affinity capture using streptavidin-conjugated beads. MWM, Molecular weight pre-stained markers. Relevant complex proteins or biological processes are indicated on the right.

In the PFDN5α enriched complexes, only four of those protein interaction networks were enriched, the prefoldin complex, the CCT/TRiC chaperonin complex, the HAUS augmin-like complex and the multiple interconnected complexes where HDAC1 was central (Figure S7A).

In the PFDN5γ enriched complexes, the prefoldin complex was no longer present, yet the chaperonin complex, the R2TP complex, the HAUS augmin-like, the 19S regulatory subunit of the 26S proteasome but also new complexes were enriched such as the Condensin I complex (Uhlmann, 2016) (Figure S7B).

Common targets of PFDN5α and PFDN5γ (intersection of PFDN5α and PFDN5γ) included the CCT/TRiC chaperonin complex, multiple PINs centered around HDAC1-RCOR1-KDM1A and quite unexpectedly to a small focal adhesion related complex composed of VCL, TLN1 and PTK2/FAK (Wehrle-Haller, 2012) (Supplementary Figure S7C).

Interestingly, in the PFDN5α specific group of protein, only the prefoldin complex was considered as enriched in the CORUM database suggesting that PFDN5α is specifically integrated in the PFD complex (Supplementary Figure S7D).

The PFDN5γ specific group of protein was enriched in CUL4A/B-DDB1 complex, an E3 ubiquitin ligase complex (Iovine et al., 2011) but also in some nuclear complexes such as the SNF2h-cohesin-NuRD complex (Hakimi et al., 2002), the Condensin complex or the 3 NELF complex (Narita et al., 2003) (Supplementary Figure S7E).

It should be noted that while the HAUS augmin-like complex is found enriched in both the PFDN5α enriched complexes and the PFDN5γ enriched complexes, it failed to be considered as enriched in the intersection of PFDN5α and PFDN5γ complexes probably because not all interaction partners are found with PFDN5α or PFDN5γ (PFDN5α 6/8 partners, PFDN5γ 5/8 partners and intersection of PFDN5α and PFDN5γ 4/8). This complex is particularly interesting as it links the known role of PFDN5 in the proper folding of the tubulins with the role of the HAUS augmin-like complex in the building of a robust mitotic spindle that is composed of such tubulins.

### 6) Identification of Novel Functional Protein Complexes Associated with both PFDN5α and PFDN5γ through Gene Ontology Enrichment Analysis of Their Proximity Interactomes

To complement the analysis performed using CORUM, we sought to identify additional proteins associated with functional complexes by employing Cytoscape with a Gene Ontology-based approach (Shannon et al., 2003).

#### 6a) PFDN5 Isoforms Associate with Diverse Chromatin Regulators

While nuclear functions of PFDN5 have been described (Blanco-Touriñán et al., 2021; Millán-Zambrano et al., 2013; Payán-Bravo et al., 2021; Pells et al., 2015), our present findings indicate that both isoforms are probably involved in these roles. Histone Deacetylase 1 (HDAC1) and Histone Deacetylase 2 (HDAC2) are conserved enzymes that form part of histone deacetylase complexes, which regulate histone deacetylation. This post-translational modification modulates chromatin accessibility and, consequently, gene expression, either activating or repressing transcription depending on the cellular context. Previous studies have identified HDAC1 and HDAC2 as protein partners of PFDN5 (Farooq et al., 2013; Joshi et al., 2013; Marcon et al., 2014). Furthermore, Satou *et al*. reported that both PFDN5α and PFDN5γ are associated with the SIN3A/TIF1β/HDAC1 complex to repress c-Myc transcriptional oncogenic activity (Satou et al., 2001). Consistently, we also detected HDAC1 and HDAC2 (Figure 6B), as well as other components of the HDAC1/2 complexes, among the proximal proteins of both PFDN5α and PFDN5γ (Supplementary table S4).

Beyond HDAC1/2, the comprehensive analysis of proteins proximal to both PFDN5α and PFDN5γ isoforms revealed a cluster of chromatin regulators involved in histone modification. We identified 32 proteins distributed across 10 distinct Gene Ontology (GO) groups (Figure 6A, Supplementary Table S4). Among these, Histone Deacetylases HDAC1 and HDAC2 were prominently detected, along with multiple components of their associated complexes (Asmamaw et al., 2024). SIN3A, a well-documented interactor of PFDN5 (Satou et al., 2001) and listed in the BioGRID database as a unique PFDN5-binding protein (see section 2b), was confirmed in our analysis. However, our results extend beyond SIN3A, revealing interactions with additional HDAC complexes: i) KDM1A, RCOR1, and RCOR3 (CoREST complex), ii) CHD3, MTA1, and MTA3 (NuRD complex), iii) MIDEAS (Asmamaw et al., 2024). Additionally, we identified TBL1XR1, a component of the N-CoR HDAC3 complex, indicating that PFDN5 proximity is not restricted to HDAC1/HDAC2 (Supplementary Table S8).

Beyond HDAC complexes, our analysis uncovered interactions with: Non-HDAC histone-modifying enzymes (e.g., KDM3A, (Yamane et al., 2006)), components of the MLL1/MLL complex (e.g., KMT2A) (Patel et al., 2009), the BRE1 complex (e.g., RNF20, RNF40) (Kim et al., 2005), the PAF1 transcription-regulating complex (e.g., LEO1) (Kim et al., 2010) and the NuA4 histone acetyltransferase complex (e.g., EP400) (Doyon and Côté, 2004)… These findings demonstrate that PFDN5 isoforms are in close proximity to a broad spectrum of histone-modifying enzymes and transcriptional regulators, extending beyond deacetylases. Collectively, our data suggest that both PFDN5α and PFDN5γ interact with diverse chromatin remodeling complexes, implicating them in epigenetic regulation (Supplementary Table S8).

To biochemically validate these observations, components of nuclear regulatory complexes, including HDAC1, HDAC2, LSD1, and SIN3A, were associated with BirA-tagged PFDN5α and PFDN5γ in BioID experiments (Figure 6B), with comparable signal intensities for both isoforms (Figure 6C).

#### 6b) Unexpected Role of PFDN5 Isoforms in Mitotic Microtubule Organization

Consistent with the identification of the HAUS augmin-like complex with the CORUM approach, a second group of proximal proteins identified by Gene Ontology with Cytoscape unexpectedly revealed an association with spindle function and microtubule organization, a cellular process not previously linked to PFDN5. This group comprised 36 proximal proteins (Figure 6A, Supplementary Table S8), including: i) four components of the octameric HAUS augmin-like complex (HAUS1,3,5, and 6), which plays a critical role in microtubule amplification during mitosis (Lawo et al., 2009; Uehara et al., 2009), ii) microtubule motor kinesins (KIF4A, 11, and 14), essential for spindle assembly and chromosome segregation (Owida et al., 2025), iii) gamma-tubulin complex proteins (TUBCG2 and 3), suggesting that a fraction of PFDN5 isoforms may localize near the centrosome (Mukhopadhyay et al., 2026), a subcellular localization not previously reported for PFDN5, iv) centrosomal proteins CEP170 (Guarguaglini et al., 2005) and CEP192 (Gomez-Ferreria et al., 2007) and the kinetochore-associated protein SKA3 (Gaitanos et al., 2009), further supporting a potential role in mitotic spindle dynamics (Supplementary Table S8).

To confirm the proximity of PFDN5 isoforms to spindle components, we focused on HAUS6, a member of the HAUS augmin-like complex and we immunoblotted BioID pull-down fractions using an anti-HAUS6 antibody. To strengthen the specificity of proteins found biotinylated by one or both PFDN5 isoforms, we included a construct expressing GFP-fused BirA (GFP-BirA) as an additional negative control alongside the BirA protein alone. We also designed truncated PFDN5 constructs to dissect the regions mediating these interactions: PFDN5α-MidDel is a construct lacking the central region containing the two β-turns required for association with other PFDN subunits in the PFD complex. This deletion allows characterization of PFD-independent interactions. PFDN5γ-MidDel and PFDN5 C-terminal constructs are truncated versions of PFDN5, retaining only the C-terminal region (Figure 7A). *In silico* predictions using AlphaFold 3.0 indicated that removal of the central region did not significantly disrupt the overall structure of PFDN5α (Supplementary Figure S8A,B) or the α-helical organization of the C-terminal region (Supplementary Figure S8C,D,E). However, simulations of the PFD complex incorporating PFDN5α-MidDel revealed a failure to properly associate with other PFD subunits (arrow, Supplementary Figure S8F; interface predicted template modelling ipTM = 0.59 vs. ipTM = 0.72 for intact PFDN5α, Figure 3A).

BioID pull-down assays confirmed that PFDN1 and PFDN3 were no longer recovered in the bound fractions of PFDN5 constructs lacking the central β-turns, consistent with their exclusion from the PFD complex (Figure 7B, Supplementary Figure S9). In contrast, HDAC1 remained associated with all constructs, indicating that its interaction with PFDN5 is mediated by the C-terminal region and independent of PFD incorporation (Figure 7B, Supplementary Figure S9).

HAUS6 was detected in the bound fractions of PFDN5α-BirA, PFDN5γ-BirA, and all truncated PFDN5 constructs (Figure 7B Supplementary Figure S9). These results demonstrate that the C-terminal region of PFDN5 isoforms is in close proximity to spindle components, independent of PFD complex incorporation.

#### 6c) PFDN5 Isoforms Associate with mRNA Metabolism Pathways

A third group of 32 distinct proteins, unexpectedly revealed an association with mRNA metabolism and post-transcriptional gene silencing, a cellular process not previously linked to PFDN5 (Figure 6A, Supplementary Table S8). Notably, this group included three components of the CCR4-NOT complex (CNOT1, CNOT3, and CNOT10), a major cellular mRNA deadenylase that regulates mRNA stability and translation (Caulier et al., 2025). The CCR4-NOT complex interacts with TNRC6B, another proximal partner of PFDN5, which is essential for miRNA-dependent translational repression and siRNA-mediated endonucleolytic cleavage of complementary mRNAs by Argonaute family proteins (Meister et al., 2005). Additionally, the group included DDX6, a protein involved in P-body formation, i.e. cytoplasmic granules that coordinate the storage and decay of translationally inactive mRNAs (Eulalio et al., 2007). The presence of EXOSC2, a subunit of the exosome complex (which mediates RNA processing and degradation), further supports the idea that PFDN5 isoforms are linked to multiple mRNA metabolism pathways (Morton et al., 2018).(Eulalio et al., 2007; Meister et al., 2005).(Morton et al., 2018). To validate these interactions, we immunoblotted BioID pull-down fractions using an anti-DDX6 antibody. This analysis confirmed that DDX6 co-purified with both PFDN5α-BirA and PFDN5γ-BirA following streptavidin affinity purification (Figure 7B). These results suggest that PFDN5 isoforms are in close proximity to key regulators of post-transcriptional gene silencing, implicating them in mRNA metabolism and RNA granule dynamics (Supplementary Table S8).

#### 6d) PFDN5 Isoforms Are in Proximity with Nuclear Pore Complex Proteins and Ran-Dependent Transport Machinery

Our proximity interactome analysis revealed that both PFDN5α and PFDN5γ isoforms associate with multiple components of the nuclear pore complex (NPC), which mediates nucleocytoplasmic trafficking. Specifically, we identified NUP107, NUP133, and NUP153 in the shared proximitome of PFDN5 isoforms. Additionally, proteins involved in Ran-dependent nucleocytoplasmic transport were also detected, including: i) RanBP2/NUP358 (an E3 SUMO-protein ligase and Ran-binding protein), ii) RanGAP1 (a Ran GTPase-activating protein), iii) UBC9 and iv) importins IMB1/KPNB1 and IMA5 (Flotho and Werner, 2012).

Among these, RanGAP1 is particularly notable due to its dual localization at the nuclear envelope during interphase and the mitotic spindle during cell division, suggesting a role in both nucleocytoplasmic transport and mitosis. RanBP2 directly binds and promotes the sumoylation of RanGAP1, a modification that regulates its subcellular localization. During mitosis, sumoylated RanGAP1 and RanBP2 are targeted to kinetochores, highlighting their role in spindle function (Joseph et al., 2002).

Although proteomic weighted scores for some NPC proteins were modest, 20 additional NPC components exhibited higher proximity scores with PFDN5 isoforms compared to the BirA control, including: NDC1, NUP37, NUP43, NUP50, NUP54, NUP58, NUP62, NUP85, NUP88, NUP93, NUP98, NUP155, NUP160, NUP188, NUP205, NUP210, NUP214, RAE1, SEC13, and SEH1.

To validate the proximity of RanGAP1 to PFDN5 isoforms, we performed BioID experiments followed by western blot analysis. RanGAP1 appears as two distinct bands on western blots: a faster-migrating band corresponding to unmodified RanGAP1 and a slower-migrating band representing its sumoylated form (SUMO*RanGAP1) (Flotho and Werner, 2012). While SUMO*RanGAP1 was barely detectable in input fractions, it was strongly enriched in the biotinylated protein fractions recovered from PFDN5α-BirA and PFDN5γ-BirA pull-downs. Notably, SUMO*RanGAP1 showed a preferential association with both PFDN5 isoforms compared to its unmodified counterpart (Figure 7B).

These results suggest that PFDN5 isoforms may participate in Ran-dependent nucleocytoplasmic shuttling, potentially regulating protein trafficking between the nucleus and cytoplasm and/or modulate the balance between nuclear pore-associated functions during interphase and spindle-associated roles during mitosis.

The preferential association with SUMO*RanGAP1 in PFDN5α proximitome further supports a potential role for PFDN5 in regulating RanGAP1 modification and localization, which could impact mitotic progression and nuclear envelope dynamics.

#### 6e) Identification of the transcription factor ZHX2 and NF-kB regulators in the proximitome of PFDN5 isoforms

In the proximitome of PFDN5 isoforms, we detected the transcription factor ZHX2, which is implicated in cell differentiation and development and acts as an oncogene in Triple Negative Breast Cancer (TNBC) and clear cell Renal Cell Carcinoma (ccRCC) (Li et al., 2022). ZHX2 was 5-fold more abundant in the PFDN5α-BirA pull-down compared to the PFDN5γ-BirA pull-down (Supplementary Table S3) and is present in the specific PFDN5α-BirA enriched set (Supplementary Table S4). This specificity was confirmed by immunoblotting, which revealed a stronger signal for ZHX2 in the PFDN5α-BirA proximitome (Figure 7B, Supplementary Figure S9). Interestingly, ZHX2 has been shown to colocalize with NF-κB at active gene promoters (Zhang et al., 2018), suggesting a potential link between PFDN5α and NF-κB signaling. Given the association between PFDN5α and ZHX2, we investigated the presence of core components of the NF-κB signaling pathway in the PFDN5α-BirA and PFDN5γ-BirA BioID pull-downs. The NF-κB complex is composed of homo- or heterodimeric assemblies of Rel-like domain-containing proteins, including RELA/p65, NF-KB1/p50, and NF-KB2/p52 (Hoesel and Schmid, 2013). Our analysis revealed that RELA/p65 and NF-KB1/p50, and to a lesser extent, NF-KB2/p52 (data not shown) were associated with both PFDN5 isoforms (Figure 7B, Supplementary Table S4).

Together with the identification of ZHX2 (and ZHX1, Supplementary Table S4), these results suggest that PFDN5 isoforms expression may modulate NF-κB signaling, potentially influencing transcriptional regulation and oncogenic pathways in specific cellular contexts.

### 7) Identification of Protein Complexes in the PFDN5*γ*-Specific Proximity Interactome

To further characterize the PFDN5γ-specific proximal interactome using Gene Ontology (GO) enrichment, we analyzed the list of PFDN5γ-BirA-specific proteins with Cytoscape. This analysis identified only one significant GO group, which included proteins associated with chromosomal maintenance and DNA repair complexes: SMC5, a component of the SMC5-SMC6 complex, and NIPBL, PDS5A, and PDS5B, which are associated with the SMC1/SMC3 cohesin complex. While this result suggests a potential link between PFDN5γ and chromosomal maintenance, the statistical significance (p-value) was insufficient to definitively conclude whether PFDN5γ is preferentially associated with the SMC5-SMC6 complex or more broadly with general DNA repair machinery. However, additional DNA repair-related proteins were identified in the PFDN5γ-specific proximitome, including DNA damage-binding protein 1 (DDB1) and CUL4A and CUL4B. The DDB1-CUL4A/CUL4B-RBX1 complex is a cullin-RING E3 ubiquitin ligase that targets proteins for proteasomal degradation following UV-induced DNA damage (Iovine et al., 2011). We confirmed a more specific association of DDB1 with PFDN5γ by western blot analysis of the BioID pull-down fractions (Figure 7B) Furthermore, we identified the PPP4C/PP4R2 serine/threonine-protein phosphatase complex in the PFDN5γ-BirA proximitome (Supplementary Figure 7B). This complex specifically dephosphorylates γ-H2AX (H2AX phosphorylated on Ser-140), a critical step in double-strand break (DSB) repair (Chowdhury et al., 2008). This finding suggests that PFDN5γ may play a role in DNA damage response pathways, particularly in the repair of DSBs.

Finally, Another interesting protein in proximity with PFDN5γ-BirA is the actin-associated Vasodilator-stimulated phosphoprotein (VASP), a components of focal adhesion (FA) structures involved in cell adhesion (Supplementary Table S4) (Wehrle-Haller, 2012)… This proximity suggests that PFDN5γ may also exert cytoplasmic function(s). PTPN11/SHP2 and ZYXIN, two partners of VASP in focal adhesion, are also found in the PFDN5γ-BirA-specific proximitome (Supplementary Table S4). Strikingly, vinculin (VCL), talin (TLN1), Breast cancer anti-estrogen resistance protein 1 (BCAR1), paxillin (PXN) are key associated factors of FA and are present in the shared proximitome (Supplementary Figure 7C) but VCL showed a three times higher score for the PFDN5γ-BirA isoform than for PFDN5α-BirA (Supplementary Table S4). Using BioID constructs, we actually demonstrated that the amount of pulled-down VCL was indeed higher in the PFDN5γ-BirA constructs and even more with the C-terminal truncated construct PFDN5-Cter (Figure 7B, Supplementary Figure S9.

## DISCUSSION

The prefoldin complex is highly conserved and plays a critical role in protein folding (Herranz-Montoya et al., 2021). Our findings demonstrate that the full-length PFDN5α isoform can functionally replace its yeast counterpart, underscoring its conserved structural and functional integrity. In contrast, the short PFDN5γ isoform fails to rescue the *pfd5Δ* yeast mutant, likely due to the absence of key structural motifs required for proper incorporation into the prefoldin complex (Siegert et al., 2000). This is consistent with our AlphaFold predictions, which suggest that PFDN5γ lacks the necessary conformational cues to stably interact with other PFDN subunits. Indeed, PFDN5γ captures less than 10% of other PFDN subunits, indicating a loose or transient association that may result in an inactive or dysfunctional complex. This inability to rescue the yeast mutant, even partially, further supports the notion that PFDN5γ does not contribute to canonical prefoldin-mediated folding.

Multiple PFDN subunits are overexpressed across a broad spectrum of tumors, often serving as biomarkers of poor prognosis in various cancer subtypes (Herranz-Montoya et al., 2021; Miyoshi et al., 2010; Shao et al., 2024; Wang et al., 2015; Yesseyeva et al., 2020). However, the role of PFDN5 in cancer remains ambiguous, with reports describing both tumor-suppressive (Hennecke et al., 2015; Satou et al., 2001; Yu et al., 2024) and tumor-promoting activities (Alldinger et al., 2005; Shao et al., 2024; Yesseyeva et al., 2020). We postulate that this duality arises from a lack of isoform-specific analysis in previous studies. Here, we reveal that the ratio of the long PFDN5α isoform to the short PFDN5γ isoform correlates with poor prognosis in HNSCC. Given that PFDN5α is the least abundant PFDN subunit across cancer subtypes (Herranz-Montoya et al., 2021), its relative enrichment may reach a threshold that triggers prefoldin-dependent oncogenic activity. This suggests that the balance between PFDN5 isoforms is a critical determinant of their functional output in cancer.

Our BioID screen identified several known PFDN5 interactors, validating our approach. These include prefoldin and CCT subunits (Gestaut et al., 2019), proteasome subunits (Kimura et al., 2007; Shahmoradi Ghahe et al., 2024), and epigenetic regulators such as SIN3 and HDAC1/2 (Satou et al., 2001). Epigenetic dysregulation is a hallmark of HNSCC progression (Castilho et al., 2017; Gaździcka et al., 2020). Among these alterations, histone acetylation and deacetylation, mediated by histone acetyltransferases (HATs) and histone deacetylases (HDACs) respectively, play a critical role in chromatin remodeling. HDAC-mediated histone deacetylation promotes chromatin compaction and transcriptional repression, contributing to chemoresistance by restricting DNA repair protein access (Lydall and Whitehall, 2005). In HNSCC, chromatin is predominantly hypoacetylated, and HDAC inhibition with trichostatin A suppresses cell proliferation while paradoxically inducing EMT (Giudice et al., 2013). Cisplatin, the standard chemotherapeutic agent for HNSCC, faces significant resistance, which is closely linked to histone acetylation status and NF-κB signaling activity (Almeida et al., 2014). These molecular insights highlight the potential of epigenetic-targeted therapies to overcome resistance and improve clinical outcomes in HNSCC. While Banks *et al*. (2018) proposed that HDAC1 and HDAC2 are substrates of the PFD/CCT folding machinery, our data reveal a more nuanced interaction: HDAC1 and HDAC2 were detected in proximity to both PFDN5α and PFDN5γ, including a truncated PFDN5α construct lacking the central β-turns required for prefoldin complex incorporation. Moreover, HDAC1 interacted with a Carboxy-terminal region shared by both isoforms, suggesting a PFD-independent association between PFDN5 and HDAC1/2. This interaction may extend beyond folding and HDAC complex assembly, potentially influencing HDAC1/2 activity in chromatin remodeling and gene regulation. This finding holds significant clinical relevance for cancer therapy, as HDAC inhibition has emerged as a highly promising therapeutic strategy, including for head and neck squamous cell carcinoma (HNSCC) (Almeida et al., 2014; Antrobus et al., 2024; Giudice et al., 2013). Moreover, comprehensive studies identifying HDAC-associated proteins have revealed interactions with all prefoldin subunits (Farooq et al., 2013; Marcon et al., 2014), while another study detected only four PFDN subunits (PFDN2, PFDN3, PFDN5, and PFDN6) in such complexes (Joshi et al., 2013). Several PFDN subunits exhibit distinct regulatory functions within the nucleus, including PFDN5 (Payán-Bravo et al., 2018). For example, specific roles for PFDN subunits have been documented in yeast and plants, where they modulate stress responses and histone remodeling. These functions appear to be PFD-independent, as evidenced by the distinct phenotypes observed in single-mutant studies (Amorim et al., 2017; Blanco-Touriñán et al., 2021; Millán-Zambrano et al., 2013). Additionally, PFDN5 has been identified in complexes lacking other PFDN subunits, such as with Rabring7 (Narita et al., 2012). Payán-Bravo *et al*. further demonstrated that both PFDN2 and PFDN5 can be chromatin-immunoprecipitated and influence a broad spectrum of gene expression (Payán-Bravo et al., 2021). The PFD-independent functions of PFDN5 are further supported by its association with chromatin-associated proteins, including components of HDAC and HAT complexes, SNF2h-cohesin-NuRD, and NELF (Doyon and Côté, 2004; Hakimi et al., 2002; Lee et al., 2022; Narita et al., 2003). While we confirmed the association of both PFDN5 isoforms with the SIN3A HDAC complex, we did not detect c-MYC in any of our BioID experiments, suggesting that c-MYC regulation by PFDN5 may be cell-type or context-specific (Fujioka et al., 2001; Satou et al., 2004, 2001). Intriguingly, PFDN1 has also been implicated in transcriptional regulation of cyclin A, although it remains unclear whether this function is PFD-dependent (Wang et al., 2017). Together, these findings point to an unexpected and broad role for PFDN5 isoforms in shaping the transcriptional landscape, independent of their canonical chaperone function.

Beyond its role in transcriptional regulation, PFDN5 is also associated with multiple components of the Ran nucleocytoplasmic transport system. Notably, PFDN5 exhibits a preferential association with the SUMOylated form of RanGAP1, suggesting a potential regulatory role in modulating the functional balance of Ran GTPase between its mitotic spindle- and interphase nuclear envelope-associated activities (Flotho and Werner, 2012). This interaction raises the possibility that PFDN5 may contribute to fine-tuning protein trafficking at the nuclear pore complex, thereby regulating the selective transport of specific proteins between the cytoplasm and the nucleus. Given that Ran has been identified as a prognostic marker in head and neck squamous cell carcinoma (HNSCC) (Zhang and Sun, 2020), these findings underscore the importance of further investigating PFDN5’s role(s) in Ran-dependent nucleocytoplasmic transport and its implications for HNSCC progression.

Our proximitome analysis also uncovered isoform-specific interactions, such as the association of PFDN5γ with DDB1 and CUL4A/B (Iovine et al., 2011), which are key players in DNA repair pathways. This suggests a potential role for PFDN5γ in genome stability, although the precise mechanisms remain to be elucidated. The prefoldin (PFD) complex has been shown to play a critical role in cell migration by mediating actin folding (Fan et al., 2020). Although PFDN5γ has thus far only been characterized in the context of its association with the HDAC-SIN3A complex (Satou et al., 2001), PFDN5γ also exhibited proximity to focal adhesion machinery components, implicating it in cell adhesion and migration, a process critical for HNSCC progression (Wehrle-Haller, 2012; Zhang and Sun, 2020). This is particularly relevant, as focal adhesions are known drivers of cancer metastasis.

The prefoldin complex is known to regulate microtubule (MT) dynamics through tubulin folding (Chesnel et al., 2020; Delgehyr et al., 2012; Rommelaere et al., 2001). However, our data reveal a striking proximity of PFDN5 to spindle-associated proteins, including γ-tubulin (TUBGCP2 and TUBGCP3) (Mukhopadhyay et al., 2026) and HAUS augmin-like complex subunits (HAUS proteins) (Lawo et al., 2009; Uehara et al., 2009), as well as MT-dependent motor proteins (KIF family) (Owida et al., 2025). Since these proteins were not identified as PFD substrates (Supplementary Table S5), their association with PFDN5 likely reflects a PFD-independent function in MT dynamics. The C-terminal region of PFDN5 appears to mediate these interactions, confirming a PFD-independent function in cell division processes. Collectively, these findings suggest that PFDN5 isoforms may interact with and influence mitotic microtubule functions, including spindle organization, a role not previously described in the literature.

This is further supported by recent evidence of PFDN5 binding to microtubules and regulating the MT-associated Tau protein aggregation both in a folding-dependent and independent manners (Bisht et al., 2026). Given that HNSCC patients are often treated with MT-targeting agents such as paclitaxel (Kitamura et al., 2020), these findings may open new avenues for understanding and targeting PFDN5 in cancer therapy.

Shao *et al*. (2004) previously demonstrated that PFDN5 expression correlates with immune response signatures and poor prognosis in gastric cancer, a finding recently confirmed in breast cancer (Wen et al., 2024). Our proximitome analysis detected multiple members of the NF-κB signaling pathway, which is known to promote tumor angiogenesis, metastasis, and resistance to chemotherapy and radiotherapy in HNSCC (Allen et al., 2007; Karin and Greten, 2005; Nakanishi and Toi, 2005). In HNSCC, the expression and activity of NF-κB is often upregulated, and its protein level increases gradually from pre-malignant lesions to invasive cancer (Bindhu et al., 2006; Ondrey et al., 1999), which suggests that NF-κB signaling plays an important role at the early stages of HNSCC carcinogenesis. A modulatory role for PFDN5 in NF-κB signaling thus warrants further investigation given the therapeutic relevance of NF-κB inhibition in HNSCC (Allen et al., 2007). We also detected a specific interaction between PFDN5α and ZHX2 (and ZHX1, Supplementary Table S4), a zinc-finger and homeobox family transcriptional regulator known to promote cell proliferation, immunoregulation, and NF-κB activity (Li et al., 2022; Zhang et al., 2018). This association further underscores the potential of PFDN5 as a modulator of immune response pathways in HNSCC, where NF-κB signaling is a critical driver of tumorigenesis and treatment resistance.

Our study provides the first comprehensive analysis of PFDN5 isoform-specific functions, revealing unexpected roles in chromatin dynamics, MT regulation, and immune signaling, independent of its canonical prefoldin chaperone activity. The proximity of PFDN5 to key regulators of spindle organization, transcriptional complexes, and NF-κB signaling highlights its potential as a novel biomarker and therapeutic target in HNSCC. Future work should focus on dissecting the mechanistic contributions of PFDN5 isoforms to these pathways, particularly in the context of HNSCC progression and treatment resistance. Given the ubiquitous nature of prefoldin-dependent folding, targeting PFDN5’s PFD-independent functions, such as its interactions with HDACs, NF-κB regulators, and MT-associated proteins, may offer a more selective therapeutic strategy to combat HNSCC while sparing normal cells.

## ACKNOWLEDGMENTS

This work was supported by a grant from La Ligue Contre le Cancer CD35 and CD29 subvention recherche 2024. We thank members of the Gene Expression and Development team at IGDR for helpful discussions. We would like to acknowledge Stéphane DREANO for his contribution in plasmid sequencing, Audrey ROUSSEL for help in biochemical experiments, Erwan WATRIN for sharing antibodies and Kevin MACE for his help with AlphaFold. We acknowledge the MRic microscopy platform at the the UAR 3480 CNRS - US18 Inserm BIOSIT.

## ABBREVIATIONS

PFD, GIMc, CCT, HNSCC, TCGA, MT, PSP (prefoldin substrate proteins)

## MATERIALS AND METHODS

### RNA extraction and RT-PCR analysis

Total RNA was extracted from cells using the Nucleospin RNA reagent kit (Macherey-Nagel). cDNAs were synthesized from 2 μg total RNA using 500 ng random hexamer primers and 200 U M-MLV reverse transcriptase (Promega) per reaction as recommended by the manufacturer. RT-PCR was performed as previously described (Chesnel et al., 2020).

### Cell culture

FaDu cell line [ATCC HTB-43] were cultivated at 37°C, 5% CO2 in DMEM medium [Gibco] with 4,5 g/L D-glucose, 4 mM L-glutamine and sodium pyruvate supplemented with 10% FBS and 1% penicillin-streptomycin.

### Yeast experiments

*S*. *pombe* strains used in this study were *h- leu1-32 ura4-D18* (WT, XLG219) and *h- pfd5::kanMX6 leu1::[nmt1-GFP-VHL213 .ura+] ura4-D18 ade6-* (XLG1048) (Chesnel et al., 2020).. Plasmids were constructed with the pADH415 backbone and contained human PFDN5α, GFP-PFDN5α and PFDN5γ. Media and genetic methods were as described (Moreno et al., 1991). Cells were grown at 25°C in Edinburgh Minimal Medium (EMM), containing with 7.5 ug/ml Thiabendazole (Sigma) when indicated. Transformation of plasmids into *S*. *pombe* strains were performed by the lithium acetate procedure. Nuclei were stained with 200 ng/ml DAPI (Sigma). Quantification of nucleus-to-cell-end distance was performed with the plot profile tool of ImageJ on inverted images of DAPI stained interphase cells (at least n = 30 cells per condition, n=3 independent experiments).

### Western blot

Proteins were resolved by SDS–PAGE and transferred onto nitrocellulose membranes. Unless otherwise specified, membranes were blocked with 5% non-fat dry milk in TBS/0.1% Tween20 (TBST/milk) and incubated overnight at 4°C with the indicated primary antibodies diluted in TBST/milk. Horseradish peroxydase-conjugated secondary antibodies used were purchased from Jackson Immunoresearch: rabbit α-mouse IgG (1:25,000), goat α-rabbit IgG (1:30,000) and goat α-rat IgG (1:10,000). Immunocomplexes were detected with ECL Select substrate (GE Healthcare) on the Amersham gel imager 680 (GE Healthcare). Quantification of Western blots by densitometry was performed using the ImageQuant software (GE Healthcare). GADPH was used as loading control to normalize the quantification.

### Co-Immunoprecipitation

ChromoTek’s GFP-Trap® is a ready-to-use reagent for the immunoprecipitation (IP) of GFP fusion proteins, and it was used in accordance with the manufacturer’s instructions.

### Subcellular fractionation

Pellets with 2.10^6^ FaDu cells expressing the protein of interest were extracted with the NE-PER extraction reagents kit (ThermoScientific) according to the manufacturer’s instructions. Subcellular fractions were quantified with the DC Protein Assay kit (Bio-Rad) and 12 ug of each fraction were loaded onto SDS-PAGE 29:1 gels for immunodetection.

### BioID

FaDu cells plated in 15-cm dishes were transfected with pCDNA3.1-BirA(R118G)-HA or pCDNA3.1-PFDN5α/γ-BirA(R118G)-HA plasmids using JetOprimusJetOptimus (Ozyme) as recommended by the manufacturer and further cultured in media supplemented with 50 μM biotin from 4 to 24 hours following transfection. They were washed 3 times in PBS, harvested and stored at -70°C until use. Pelleted cells were lysed in RIPA buffer (50 mM Tris-HCl, pH 7.6, 150 mM NaCl, 1 mM EDTA, 1 mM EGTA, 0.1% SDS, 1% Triton X100 and protease inhibitor cocktail (Thermo Fisher Scientific)), sonicated and centrifuged at 16,000 g for 15 min at 4°C. Total proteins were quantified by Bio-Rad assay and equal amounts of total proteins in each experiment were used for streptavidin capture. Before the affinity chromatography, aliquots of total proteins were analyzed for each condition (“input” fractions). Streptavidin-Sepharose Beads (60 μl, BioVision) were washed 3 times with lysis buffer and incubated with total protein lysates for 3h30 at 4°C on a tube rotator. Protein-bound beads were then washed three times with RIPA buffer and three times with 50 mM ammonium bicarbonate pH 7.8. Elution of biotinylated proteins (“Bound” fraction) was carried out by adding 75 μl 2xLaemmli loading buffer to the beads and eluted proteins were denatured by heat at 95°C for 5 min prior to western blot analysis.

### Immunofluorescence

FaDu cells were cultivated on 18 mm-glass coverslips for 24 h beforetransfection with peGFP-C1-PFDN5α/γ or pCDNA3.1-PFDN5α/γ-HA plasmids using JetOptimus (Ozyme) as recommended by the manufacturer then cultivated for 24-30 hours following transfection. Cells werefixed with 4% paraformaldehyde for 10 min, permeabilized with 0.2% Triton X-100 in PBS for 5 min and stained after blocking 1 hour with 1% BSA in PBS. Primary anti-GFP (mouse clones13.1 & 7.1, Roche) or anti-HAtags antibodiesrat clone 3F10, Roche) at dilution 1:100), were incubated onto the cells for 2 h at room temperature. overnight at 4°C.After washing, the dye conjugated-secondary antibodies directed against mouse or rat IgG (Jackson ImmunoResearch, dilution 1:1000) were incubated for 1h at room temperature. After washing with PBS/0,1%. Tween20, the nuclei were stained with DAPI and-coverslips were mounted with Prolong Gold^TM^Antifade (Invitrogen). Cells were captured using Confocal Leica DMI 6000 SP8 Inverted Microscope with the Leica software.

### Mass spectrometry proteomics analysis

Tryspin digestion was carried out as previously described with minor modifications (Sandoval Pacheco et al., 2025). Briefly beads were centrifuged 3 min at 320 x g before resuspended in 20 µL of UTH-Pmax Buffer (6M Urea, 50 mM Tris-HCl pH8, 0,01% ProteaseMAX^TM^ (Promega)). Proteins were then subjected to reduction using 7,2 mM DTT in UTH-PMax buffer (see above) for 15 minutes at 37 °C. Alkylation was then performed using 13,5 mM iodoacetamide in UTH-PMax buffer (see above) for 15 minutes at room temperature in the darkness. The samples were then digested with 0,4 µg of a Tryspin/Lys-C mix (V507A, Promega) at 37 °C for 3h. The digestion was then continued overnight after adding 50mM Tris-HCl pH8 and 0,01% ProteaseMAX^TM^ to reach 4ng/µL of enzyme. The resulting peptide mixtures were then cleaned up from salts, contaminants, and detergents using Phoenix cartridges (PreOmics GmbH, Martinsried, Germany).

Approximately 350 ng each of tryptic peptides samples were separated onto a 75 μm × 250 mm IonOpticks Aurora 3 column (Ion Opticks Pty Ltd., Australia) packed with a 120 Å pore, 1.7 μm particle size C18 beads. A reversed-phase gradient of basic buffers (buffer A: 0.1% formic acid, 98% H2O Milli-Q, 2% acetonitrile; buffer B: 0.1% formic acid, 100% acetonitrile) was run on a NanoElute high-performance liquid chromatography (HPLC) system (Bruker Daltonik) at a flow rate of 250 nL/min at 50 °C. The LC run lasted for 80 min with a starting concentration of 2% buffer B increasing to 13% over the first 42 minutes was first performed and buffer B concentrations were increased up to 20% at 65 min; 30% at 70 min; 85% at 75 min and finally 85% for 5 min to wash the column. The NanoElute HPLC system was coupled online to a Tims time-of-flight (TOF) Pro mass spectrometer (Bruker Daltonik) with a CaptiveSpray ion source (Bruker Daltonik). The CaptiveSpray nanoflow electrospray (ESI) source was directly attached to a vacuum inlet capillary via a short capillary extension heated using the instrument’s drying gas. High voltage for the ESI process was applied to the vacuum capillary inlet, whereas the sprayer was kept at ground. The temperature of the ion transfer capillary was set at 180°C. The spray type was automatically mechanically aligned on the axis with the capillary inlet without the need for any adjustment. Ions were accumulated for 100ms, and mobility separation was achieved by ramping the entrance potential from −160 to −20 V within 114 ms. The acquisition of the MS mass spectra with the trapped ion mobility spectrometry (TIMS) TOF Pro is done with an average resolution of 60 000 fwhm (mass range 100− 1700 m/z). To enable the PASEF method, precursor m/z and mobility information was first derived from full scan TIMS-MS experiments, MS and MS/MS data were collected over the m/z range 254.1 - 1200 and over the mobility range from 1/K0 = 0.75 to 1/K0 = 1.33 Vs cm-2. Resulting quadrupole mass, collision energy and switching times were automatically transferred to the instrument controller as a function of the total cycle time. The quadrupole isolation width was set to 2 and 3 Th and, for fragmentation, the collision energies varied between 20 and 59 eV depending on precursor mass and charge. TIMS, MS operation and PASEF were controlled and synchronized using the control instrument software timsControl 6.0 (Bruker Daltonik). LC-MS/MS data were acquired using the PASEF method with a total cycle time of 1.17 s, including 1 TIMS MS scan and 10 PASEF MS/MS scans. The 10 PASEF scans (100 ms each) contain on average 20 MS/MS scans per PASEF scan. In addition, the most abundant precursors which could have been sequenced in previous scan cycles are dynamically excluded from resequencing. The acquisition of the MS/MS mass spectra with the TIMS TOF Pro is also done with an average resolution of 50 000 fwhm (mass range 100−1700 m/z). Ion mobility resolved mass spectra, nested ion mobility vs m/Z distributions, as well as summed fragment ion intensities were extracted from the raw data file with DataAnalysis 6.0 (Bruker Daltonik). Signal-to-noise (S/N) ratios were increased by summations of individual TIMS scans. Mobility peak positions and peak half-widths were determined based on extracted ion mobilograms (±0.05 Da) using the peak detection algorithm implemented in the DataAnalysis software. Feature detection was also performed using DataAnalysis 6.0 software and exported in .mgf format.

Peptide and protein identification were performed using the Mascot (Mascot server v2.6.2; http://www.matrixscience.com) database search engine. MS/MS spectra were queried against the UniProtKB Homo Sapiens proteome database UP000005640 UniProtKB (release December 2022) restricted to one protein sequence per gene (20594 sequences), completed with the sequences of the BirA-HA, the Pfdn5a-BirA-HA, the Pfdn5g-BirA-HA and a common proteomic contaminant database from the Max Planck Institute of Biochemistry, Martinsried (247 sequences). Mass tolerance for MS and MS/MS was set at 15 ppm and 0.05 Da. The enzyme selectivity was set to trypsin with one miscleavage being permitted. The peptides modifications that have been considered were carbamidomethylation of cysteines as fixed modifications, and the oxidation of methionine, the acetylation of lysine, the acetylation of N-terminal proteins and the biotinylation at any N-terminal of peptides as variable modifications. Identification results from Mascot (.dat files) were imported into the Proline Studio software (Bouyssié et al., 2020). This software was then used to validate protein identification with a peptide rank = 1 and a 0,1% FDR based on the Benjamini-Hochberg procedure at the peptide spectrum matches level (Couté et al., 2020). Proteins identified with exactly the same set of peptides or with a subset of the same peptides were grouped in a Protein Set which is represented by the best-identified protein (best score) or in case of same set proteins, the one with a SwissProt accession if possible. When proteins with shared peptides were identified with other peptides not belonging to the Protein Set, different Protein Sets were created, even if there are no specific peptides (i.e., if theses peptides were also shared by other Protein Sets). Proline Studio software was also used to the spectral count comparison of the identified proteins in each sample as previously described (Simoes Eugénio et al., 2021). For each protein, a weighted spectral count is calculated, as suggested in Abacus (Fermin et al., 2011), where shared peptides are combined and weighted according to the specific spectral counts of the different Protein Sets sharing the same peptide(s). To detect significant difference between samples, a beta-binomial test was performed on these weighed spectral counts and a p-value was calculated for each Protein Set using the R package BetaBinomial 1.2 implemented in Proline Studio (Bouyssié et al., 2020; Pham et al., 2010).(Couté et al., 2020; Simoes Eugénio et al., 2021).(Fermin et al., 2011; Pham et al., 2010).

### Mass Spectrometry Data Analysis

Gene symbols unique to PFDN5-BirA isoform sets were submitted as a gene list to CORUM online functional annotation tool (Tsitsiridis et al., 2023) and all enriched gene ontology (GO) annotations were retrieved using the Cytoscape online tool (Shannon et al., 2003). The most significant terms associated with each GO categories were selected.

### Statistics

All statistical analyses were done with R studio. The error bars represents ±S.D (unless otherwise specified) from experimental independent replicates of at least n = 3. Statistical analyses were conducted with Student t-test, Kruskal-Wallis or Mann-Whitney tests. The Kaplan-Meier survival curve was done with R using the Survival package, version 3.5-8 (Therneau TM. A Package for Survival Analysis in R. 2020. Available from: https://CRAN.R-project.org/package=survival). We used the TCGA splicing data from TCGASpliceSeq (downloaded: March, 2023) (Ryan et al., 2016).

## SUPPLEMENTARY FIGURE LEGENDS

**Supplementary Figure S1: Prognostic Impact of PFDN Subunits in HNSCC.** Kaplan-Meier survival curves for HNSCC patients based on PFDN1, PFDN3/VBP1, and PFDN4 expression (poor prognosis when expression is high), and total PFDN5 expression (no significant impact, p=0.1).

Supplementary Figure S2: Identification of BioGrid interacting proteins and putative Prefoldin Substrate Proteins in the BioID datasets. A circular chord diagram depicting the relationships among prefoldin subunits (PFDN1 to PFDN6) and their interacting proteins. Each prefoldin subunit is represented by a distinct color arc on the outer circle, with connecting chords indicating interactions with indicated proteins. Proteins associated with each subunit are labeled around the perimeter.

**Supplementary Figure S3: PFDN5*α*, but not PFDN5*γ*, associates with other prefoldin subunits in A253 HNSCC cells.** Immunoblot analysis of HA-tagged BirA alone or fused to the indicated PFDN5 isoforms and PFDN subunits, shown before (Input, left) and after (Bound, right) affinity capture using streptavidin-conjugated beads. Ctrl, untransfected control cells; MWM, Molecular weight pre-stained markers.

**Supplementary Figure S4: Representative images of fission yeast cells used to measure the nucleus-to-cell-end distance ratio.** Wild type (WT) or *pfd5*Δ cells transfected with control empty vector (vector) or plasmids allowing the production of either human PFDN5α, GFP-PFDN5α or PFDN5γ were stained with DAPI. Red arrows shown on inverted images are corresponding to exact half cells, thus indicating the geometric cell center where the arrowheads meet.

**Supplementary Figure S5: Localization of PFDN5*α*-HA isoform in FaDu cells assessed by indirect immunofluorescence.** Representative confocal microscopy images of indirect immunofluorescence of FaDu cells transfected with PFDN5α-HA, fixed and stained with an anti-HA antibody (green) and DAPI (nuclei, blue).

Supplementary Figure S6: Expression levels of recombinant HA-tagged PFDN5 isoforms driven by pPGK or pCMV promoters in FaDu cells. Immunoblot analysis of HA-tagged PFDN5 isoforms expressed under the control of either the low-expression pPGK promoter or the high-expression pCMV promoter in transfected FaDu cells. α-tubulin was used as a loading control. Ctrl, untransfected control cells. MWM, Molecular weight pre-stained markers.

**Supplementary Figure S7: Identification of PFDN5 isoform-interacting proteins in FaDu cells using BioID and LC-MS/MS.** A) All proteins identified in the most represented CORUM categories with PFDN5α (all_alpha), B) All proteins identified in the most represented CORUM categories with PFDN5γ (all_gamma), C) Common targets of PFDN5α and PFDN5γ (intersection of PFDN5α and PFDN5γ) (alpha_&_gamma). D) Specific proteins identified in the most represented CORUM categories with PFDN5α (alpha_spe), E) Specific proteins identified in the most represented CORUM categories with PFDN5γ (gamma_spe). Six main protein interaction networks (PINs) are indicated by a color code.

**Supplementary Figure S8: *In silico* AlphaFold predictions of PFDN5 structures and PFD complex formation.** A–E) Predicted structures of BirA-HA-tagged PFDN5 proteins, including full-length and truncated recombinant variants. F) AlphaFold simulation of PFD complex formation incorporating PFDN5α-MidDel-BirA-HA and the five other PFDN subunits.

Supplementary Figure S9: Quantification of biotinylated target proteins associated with PFDN5 isoforms and their truncated versions (MidDel, Cter), normalized to the protein with maximal association set as 100%. BirA and GFP-BirA were used as controls.

## SUPPLEMENTARY TABLES

Supplementary Table S1: Relative abundance of PFDN5*α* and PFDN5*γ* isoforms Supplementary Table S2: BioID samples used in this study

Supplementary Table S3: Proteomic data input and binomial test for PFDN5*α* vs BirA and PFDN5*γ vs* BirA

Supplementary Table S4: Intersection of proteomic PFDN5 targets present in both PFDN5*α* and PFDN5*γ* BioID (inBoth), only in PFDN5*α* BioID (inAonly) or only in PFDN5*γ* BioID (inGonly)

Supplementary Table S5: Matrix of Biogrid PFDNs results

Supplementary Table S6: Contingency Table for BioGrid enrichment sets for PFDN5*α*

Supplementary Table S7: Contingency Table for BioGrid enrichment sets for PFDN5*γ*

Supplementary Table S8: Proteomic PFDN5 targets present in both PFDN5*α* and PFDN5*γ* BioID and identified by a Gene Ontology-based approach with Cytoscape. Three groups based on Gene-Ontology keyword (Histone modification, Spindle/centrosome and Post-transcriptional gene silencing) are reported.

