## Supplementary figures and images for "Differential Roles of PFDN5 Isoforms in Head and Neck Squamous Cell Carcinoma: Insights from Proximity Interactome Mapping"

### Supplementary figures

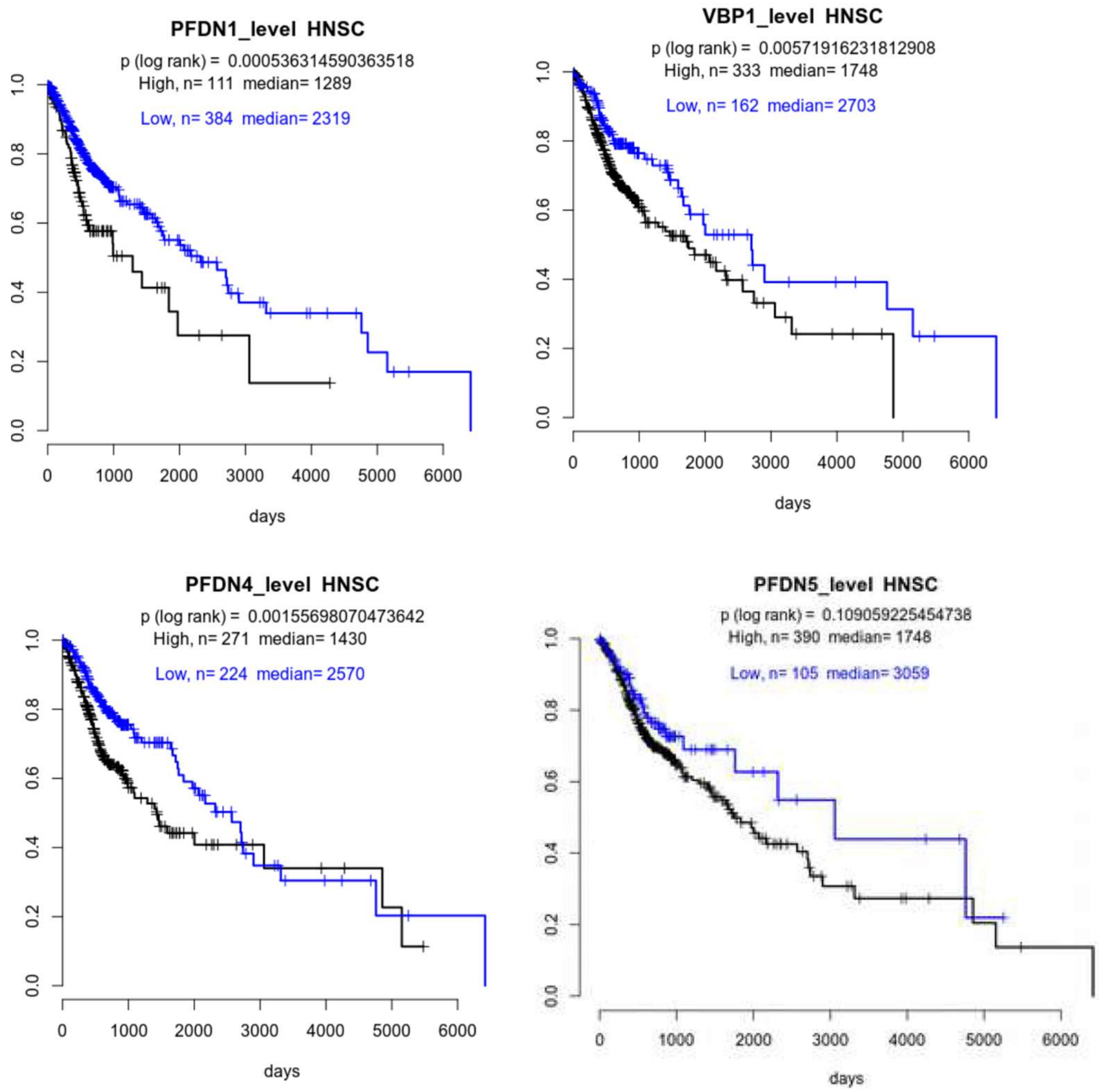

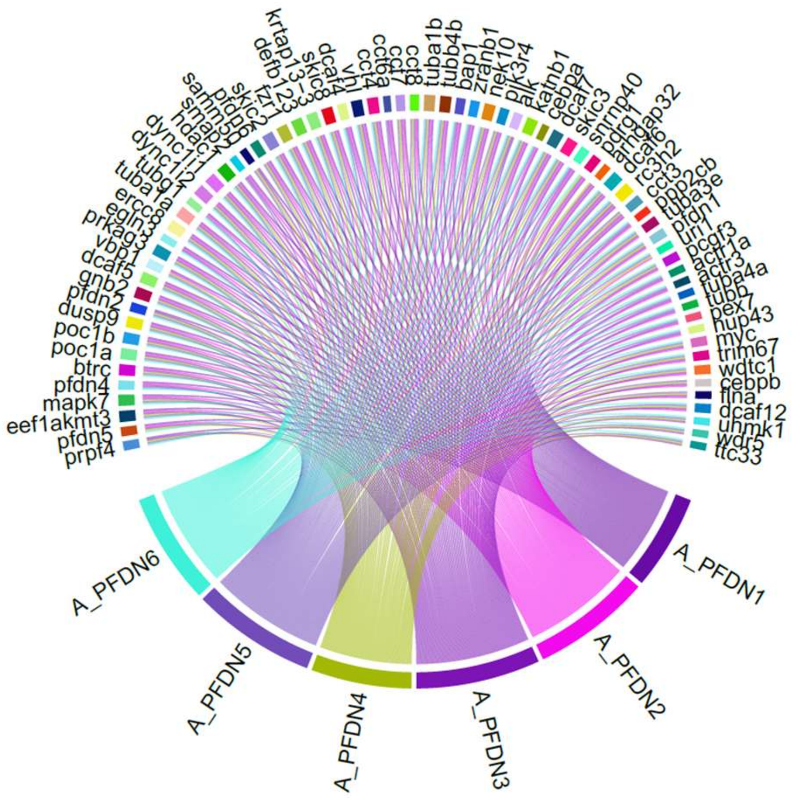


Figure S2

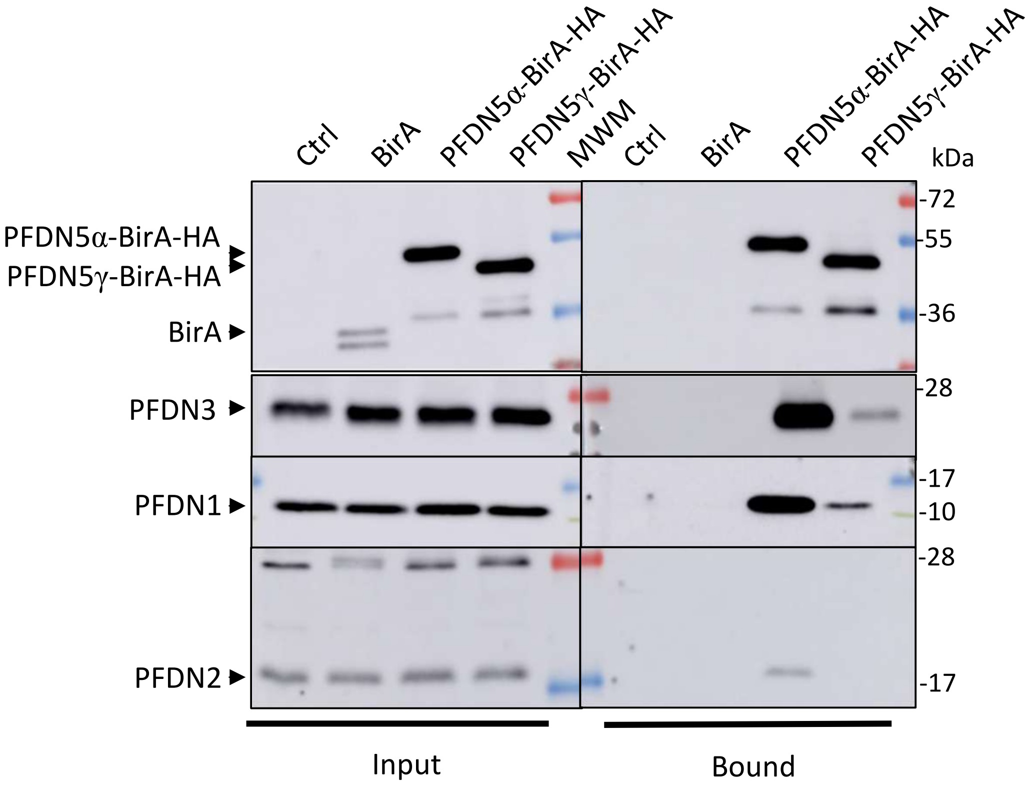

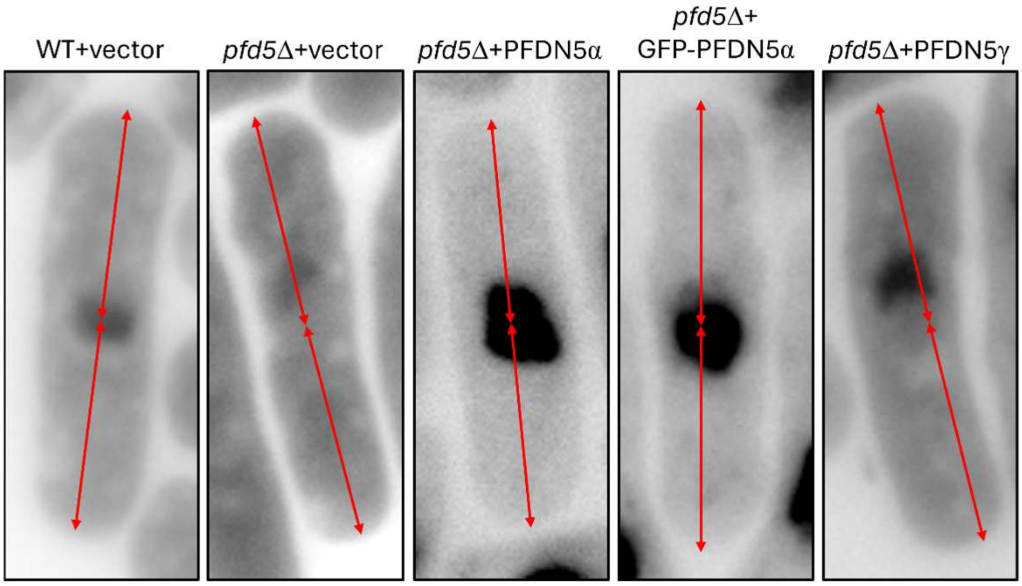

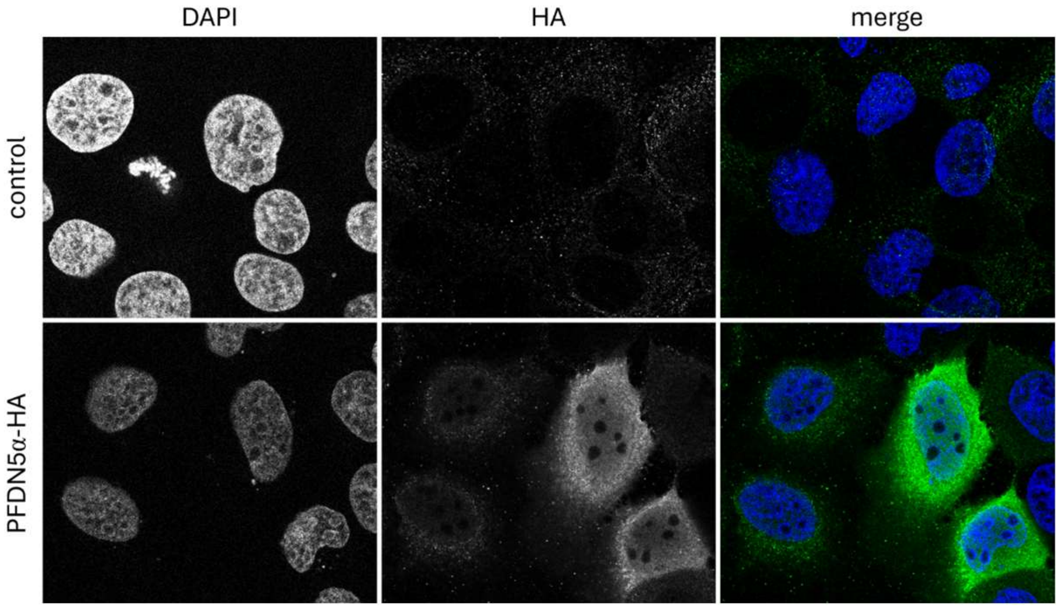


Figure S5

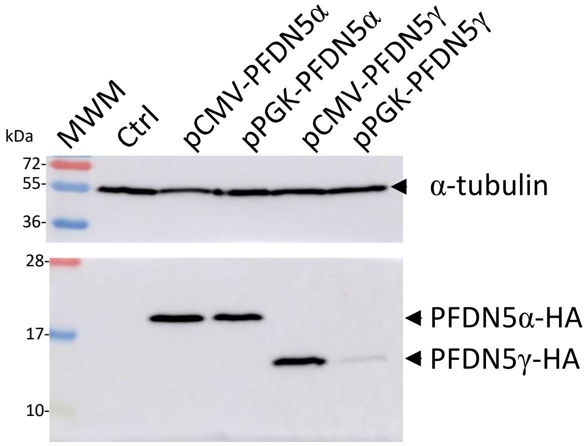

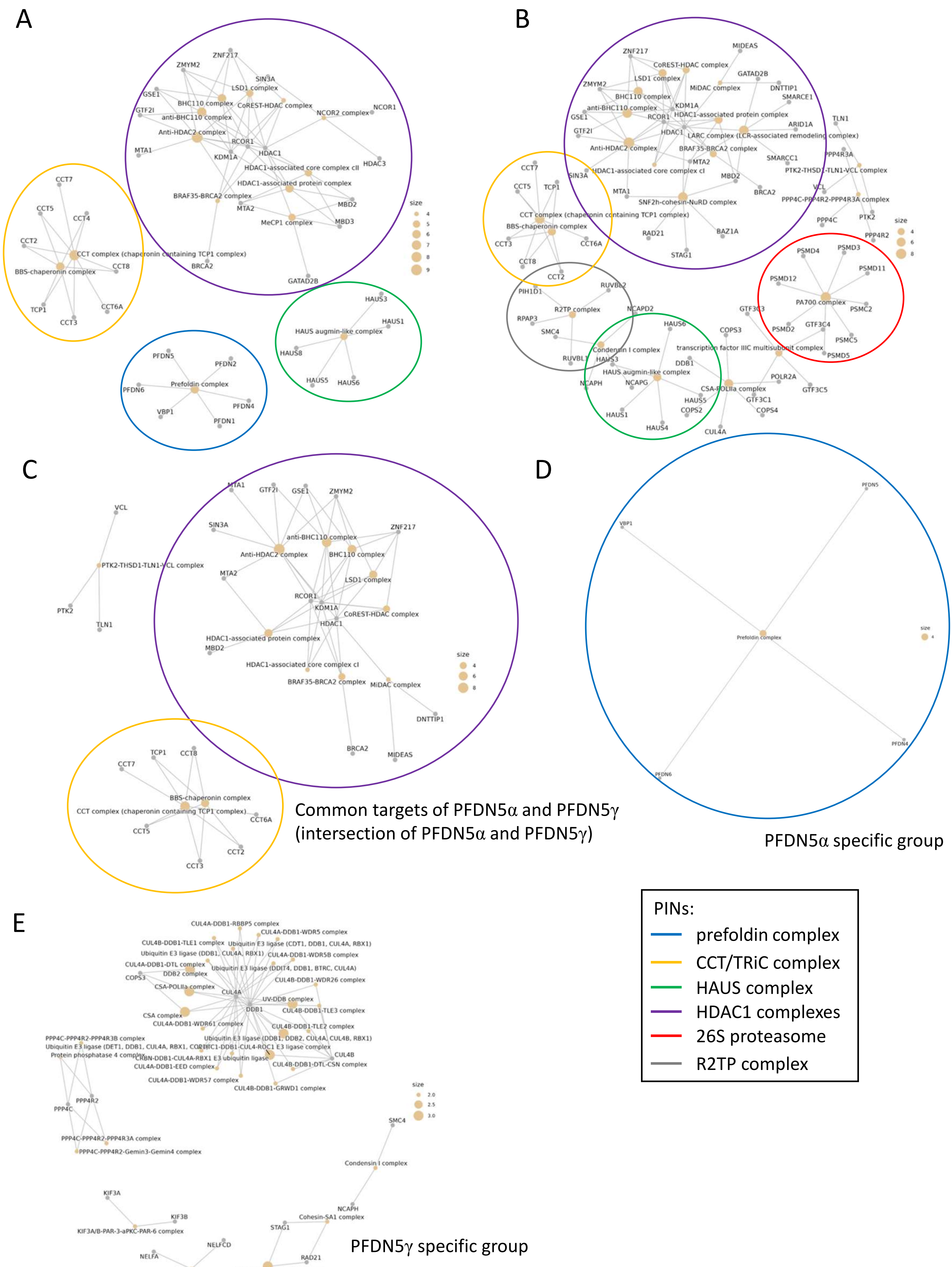

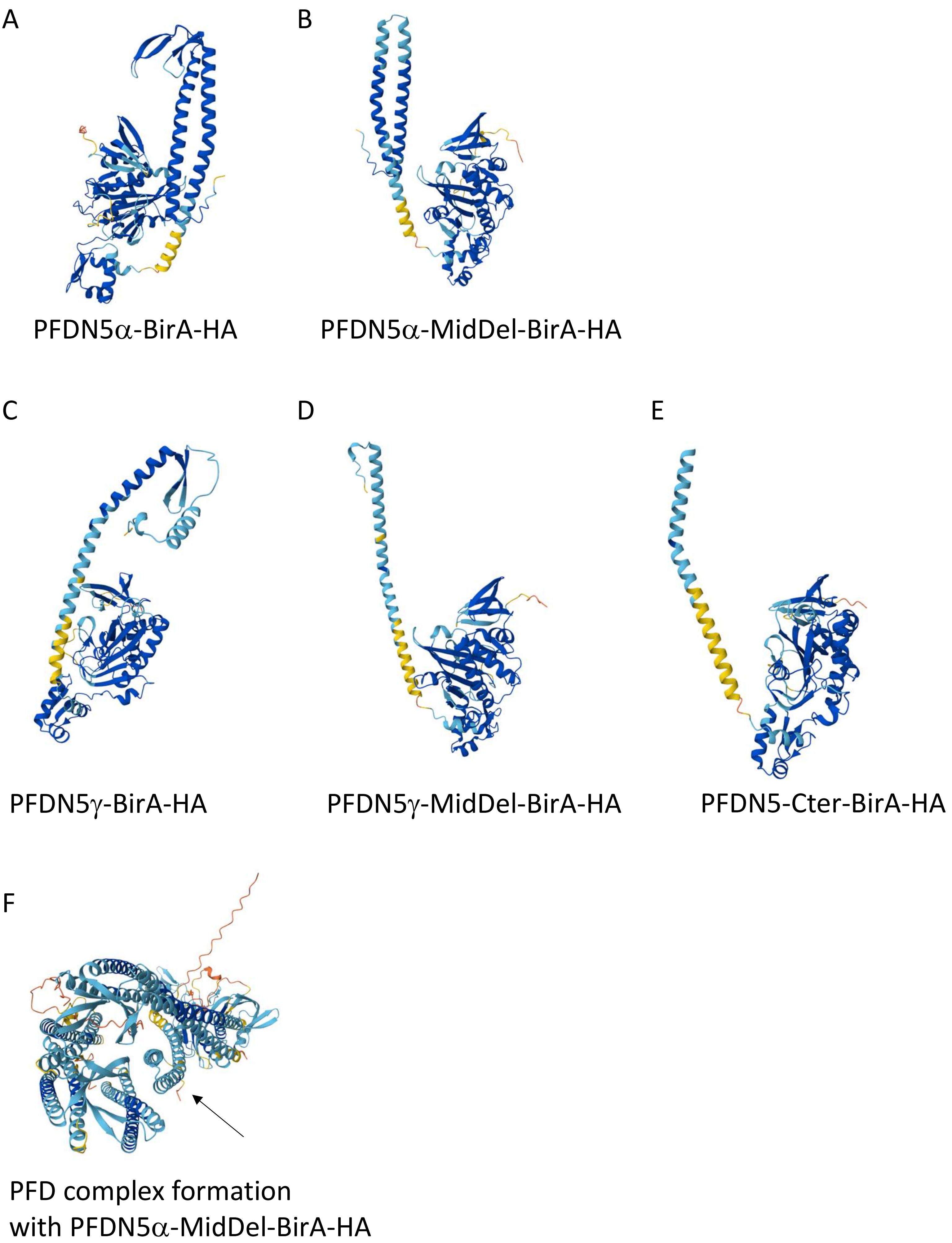

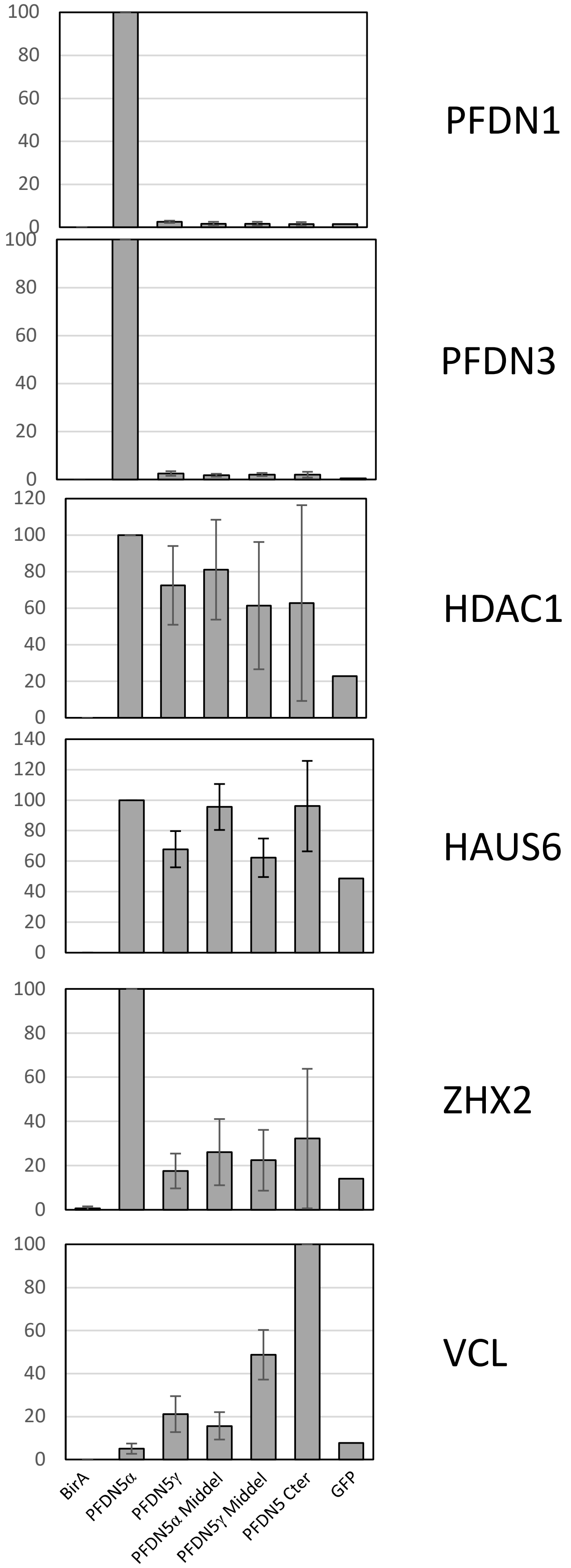
