## supplementary Table S1 for "Differential Roles of PFDN5 Isoforms in Head and Neck Squamous Cell Carcinoma: Insights from Proximity Interactome Mapping"

**Supplementary Table S1: Relative abundance of *PFDN5α* and *PFDN5γ* isoforms**

Data extracted from GTEX (TPM), except for HaCaT where the data originate from a poly(A)+ Rnaseq using Oxford nanopore long read sequencing and are reads/ junction for junction exon1-Exon 2 and junction Exon1-Exon 4

| *PFDN5α* | *PFDN5γ* | Tissue | *PFDN5α* / *PFDN5γ* |
| --- | --- | --- | --- |
| 343 | 23.2 | E-Mucosa | 14.784483 |
| 406 | 37.6 | Minor salivary gland | 10.797872 |
| 500 | 54.7 | E-Muscularis | 9.140768 |
| 547 | 56.5 | E-Gastroesophageal junction | 9.681416 |
| 329 | 129.0 | testis | 2.550388 |
| 765 | 33.0 | HaCaT | 23.181818 |
