## supplementary Table S2 for "Differential Roles of PFDN5 Isoforms in Head and Neck Squamous Cell Carcinoma: Insights from Proximity Interactome Mapping"

**Supplementary Table S2: BioID samples used in this study**

| ID | condition | Experiment | short_id |
| --- | --- | --- | --- |
| 412_T2_01_Slot2-37_1_5992.d | JP | 1 | JP_1 |
| 412_T2_02_Slot2-38_1_5993.d | BirA | 1 | BirA_1 |
| 412_T2_03_Slot2-39_1_5994.d | Gamma | 1 | Gamma_1 |
| 412_T2_04_Slot2-40_1_5995.d | Alpha | 1 | Alpha_1 |
| 412_05_Slot1-10_1_6977.d | JP | 2 | JP_2 |
| 412_06_Slot1-11_1_6978.d | BirA | 2 | BirA_2 |
| 412_08_Slot1-13_1_6980.d | Gamma | 2 | Gamma_2 |
| 412_09_Slot1-14_1_6981.d | JP | 3 | JP_3 |
| 412_07_Slot1-12_1_6979.d | Alpha | 2 | Alpha_2 |
| 412_10_Slot1-15_1_6982.d | BirA | 3 | BirA_3 |
| 412_11_Slot1-16_1_6983.d | Alpha | 3 | Alpha_3 |
| 412_12_Slot1-17_1_6984.d | Gamma | 3 | Gamma_3 |
