## supplementary Table S6 for "Differential Roles of PFDN5 Isoforms in Head and Neck Squamous Cell Carcinoma: Insights from Proximity Interactome Mapping"

**Supplementary Table S6: Contingency Table of PFDN5α interactors**

| Condition | bioID_YES | bioID_NO |
| --- | --- | --- |
| biogrid_YES | 29/15.5 | 117/130.5 |
| biogrid_NO | 540/553.5 | 4657/4643.5 |
