## supplementary Table S7 for "Differential Roles of PFDN5 Isoforms in Head and Neck Squamous Cell Carcinoma: Insights from Proximity Interactome Mapping"

**Supplementary Table S7: Contingency Table of PFDN5γ interactors**

| Condition | bioID_YES | bioID_NO |
| --- | --- | --- |
| biogrid_YES | 27/14.7 | 119/131.3 |
| biogrid_NO | 515/527.3 | 4711/4698.7 |
