## supplementary Table S8 for "Differential Roles of PFDN5 Isoforms in Head and Neck Squamous Cell Carcinoma: Insights from Proximity Interactome Mapping"

### Histone modification

| | Occurrences in cytoscape (10 groups) | p.adj.gamma | PFDN5g-BirA | p.adj.alpha | PFDN5a-BirA |
|---|---|---|---|---|---|
| BRCA2 | 2 | 0,032 | 7,7 | 0,005 | 17,3 |
| DTX3L | 2 | 0,031 | 5,0 | 0,018 | 5,0 |
| EP400 | 3 | 0,029 | 19,7 | 0,005 | 24,7 |
| GTF3C4 | 3 | 0,013 | 13,0 | 0,011 | 11,3 |
| HCFC1 | 3 | 0,009 | 30,3 | 0,005 | 54,3 |
| ★ HDAC1 | 4 | 0,041 | 8,5 | 0,017 | 11,8 |
| ★ HDAC2 | 4 | 0,031 | 12,8 | 0,014 | 15,7 |
| ➡ KDM1A | 4 | 0,008 | 16,3 | 0,005 | 20,7 |
| KDM3A | 1 | 0,022 | 4,5 | 0,006 | 9,0 |
| KMT2A | 4 | 0,008 | 11,7 | 0,031 | 14,0 |
| LEO1 | 2 | 0,014 | 5,7 | 0,044 | 6,7 |
| MAP3K7 | 4 | 0,014 | 5,3 | 0,006 | 7,3 |
| MCM3AP | 4 | 0,009 | 8,7 | 0,016 | 10,0 |
| MIDEAS | 2 | 0,015 | 14,7 | 0,005 | 22,0 |
| MTA1 | 2 | 0,012 | 13,0 | 0,005 | 15,6 |
| MTA2 | 2 | 0,025 | 15,5 | 0,009 | 19,8 |
| OGT | 4 | 0,012 | 16,3 | 0,006 | 25,0 |
| ★ PRKAA1 | 3 | 0,026 | 4,3 | 0,013 | 5,0 |
| RCOR1 | 2 | 0,014 | 5,7 | 0,041 | 5,7 |
| RCOR3 | 2 | 0,021 | 4,6 | 0,028 | 4,3 |
| RIF1 | 1 | 0,007 | 33,3 | 0,010 | 25,0 |
| RNF20 | 5 | 0,007 | 9,5 | 0,020 | 4,3 |
| RNF40 | 5 | 0,007 | 12,2 | 0,006 | 9,7 |
| ★ RUVBL1 | 3 | 0,040 | 166,0 | 0,034 | 138,7 |
| ★ RUVBL2 | 3 | 0,043 | 194,0 | 0,029 | 181,7 |
| SART3 | 1 | 0,010 | 27,3 | 0,007 | 24,7 |
| ➡ SIN3A | 6 | 0,009 | 14,3 | 0,007 | 17,7 |
| SIRT1 | 6 | 0,021 | 7,7 | 0,009 | 9,3 |
| TBL1XR1 | 2 | 0,007 | 18,3 | 0,006 | 28,0 |
| UBR5 | 3 | 0,024 | 13,7 | 0,005 | 24,0 |
| WDR70 | 4 | 0,013 | 10,0 | 0,012 | 8,3 |
| YEATS2 | 3 | 0,008 | 8,7 | 0,005 | 17,7 |

| | |
|---|---|
| (yellow) | alpha = ~2* gamma |
| (orange) | gamma = ~2* alpha |

★ Proteins found with other PFDNs in Biogrid

➡ Proteins exclusive to PFDN5 in Biogrid

### Spindle/centrosome

| | Occurrences in cytoscape (10 groups) | p.adj.gamma | PFDN5g-BirA | p.adj.alpha | PFDN5a-BirA |
|---|---|---|---|---|---|
| APC | 2 | 0,008 | 10,7 | 0,005 | 16,7 |
| ARHGEF7 | 5 | 0,010 | 7,7 | 0,021 | 6,3 |
| BRCA2 | 2 | 0,032 | 7,7 | 0,005 | 17,3 |
| CAMSAP1 | 4 | 0,013 | 19,0 | 0,005 | 19,6 |
| CAMSAP2 | 4 | 0,008 | 15,0 | 0,005 | 17,6 |
| CEP192 | 7 | 0,008 | 9,0 | 0,005 | 17,7 |
| CHD3 | 3 | 0,012 | 11,0 | 0,017 | 11,3 |
| CHORDC1 | 2 | 0,020 | 8,0 | 0,011 | 7,0 |
| CKAP5 | 8 | 0,007 | 37,0 | 0,005 | 39,3 |
| CLIP1 | 4 | 0,035 | 9,2 | 0,023 | 5,9 |
| CYLD | 1 | 0,041 | 4,3 | 0,032 | 3,7 |
| DCTN1 | 9 | 0,035 | 8,3 | 0,022 | 8,3 |
| DLG1 | 1 | 0,015 | 12,0 | 0,047 | 9,0 |
| GIT1 | 5 | 0,023 | 4,8 | 0,019 | 6,3 |
| GOLGA2 | 7 | 0,008 | 17,3 | 0,007 | 14,3 |
| HAUS1 | 4 | 0,036 | 3,7 | 0,023 | 4,0 |
| HAUS3 | 4 | 0,037 | 5,3 | 0,013 | 7,3 |
| HAUS5 | 4 | 0,015 | 5,0 | 0,011 | 5,7 |
| HAUS6 | 4 | 0,007 | 13,7 | 0,005 | 16,0 |
| KIF11 | 4 | 0,048 | 9,0 | 0,006 | 12,3 |
| KIF4A | 2 | 0,015 | 10,9 | 0,030 | 8,3 |
| KPNB1 | 3 | 0,041 | 41,7 | 0,008 | 51,7 |
| MAP4 | 1 | 0,033 | 25,7 | 0,050 | 24,7 |
| MAP7D3 | 2 | 0,007 | 9,7 | 0,005 | 9,0 |
| MPDZ | 2 | 0,025 | 5,7 | 0,013 | 6,8 |
| NCOR1 | 2 | 0,007 | 26,4 | 0,005 | 44,8 |
| PATJ | 2 | 0,010 | 11,0 | 0,006 | 12,2 |
| ★ PRKAA1 | 1 | 0,026 | 4,3 | 0,013 | 5,0 |
| SIRT1 | 2 | 0,021 | 7,7 | 0,009 | 9,3 |
| SKA3 | 2 | 0,011 | 6,3 | 0,005 | 13,0 |
| SLK | 1 | 0,008 | 19,2 | 0,009 | 16,3 |
| SPAG5 | 3 | 0,007 | 13,3 | 0,005 | 15,0 |
| TACC1 | 1 | 0,027 | 4,3 | 0,009 | 8,3 |
| TUBCGP2 | 6 | 0,012 | 6,0 | 0,023 | 4,3 |
| TUBCGP3 | 6 | 0,025 | 4,3 | 0,026 | 6,0 |
| WASHC5 | 2 | 0,007 | 11,0 | 0,005 | 10,0 |

### Post-transcriptional gene silencing

| | Occurrences in cytoscape (17 groups) | p.adj.gamma | PFDN5g-BirA | p.adj.alpha | PFDN5a-BirA |
|---|---|---|---|---|---|
| CNOT1 | 12 | 0,012 | 15,0 | 0,011 | 11,0 |
| CNOT10 | 3 | 0,010 | 8,7 | 0,012 | 6,3 |
| CNOT3 | 9 | 0,028 | 4,0 | 0,011 | 6,7 |
| CSDE1 | 2 | 0,007 | 35,3 | 0,005 | 31,7 |
| DCP1A | 6 | 0,011 | 13,7 | 0,011 | 12,7 |
| DDX6 | 11 | 0,009 | 18,7 | 0,005 | 24,3 |
| EDC4 | 1 | 0,007 | 22,0 | 0,005 | 20,7 |
| EGFR | 8 | 0,012 | 29,0 | 0,015 | 29,0 |
| EIF4E | 1 | 0,023 | 12,3 | 0,020 | 13,3 |
| EIF4ENIF1 | 17 | 0,008 | 8,7 | 0,005 | 15,7 |
| EIF4G1 | 11 | 0,016 | 30,4 | 0,011 | 32,5 |
| EXOSC2 | 7 | 0,037 | 3,7 | 0,018 | 4,7 |
| ★ HDAC1 | 1 | 0,041 | 8,5 | 0,017 | 11,8 |
| LARP1 | 2 | 0,029 | 15,3 | 0,021 | 15,3 |
| MBD2 | 1 | 0,044 | 5,3 | 0,040 | 5,3 |
| NCOR1 | 7 | 0,007 | 26,4 | 0,005 | 44,8 |
| NCOR2 | 7 | 0,007 | 15,6 | 0,005 | 35,8 |
| PATL1 | 7 | 0,036 | 3,7 | 0,006 | 8,3 |
| RBM10 | 1 | 0,009 | 17,6 | 0,006 | 17,2 |
| ★ RC3H2 | 7 | 0,043 | 3,5 | 0,006 | 11,3 |
| RESF1 | 2 | 0,020 | 5,0 | 0,023 | 4,0 |
| SIRT1 | 1 | 0,021 | 7,7 | 0,009 | 9,3 |
| SMG1 | 1 | 0,018 | 5,0 | 0,020 | 7,0 |
| SMG7 | 1 | 0,011 | 8,7 | 0,005 | 11,7 |
| STAT3 | 11 | 0,036 | 29,3 | 0,026 | 22,0 |
| TASOR | 2 | 0,009 | 20,0 | 0,005 | 16,7 |
| TNRC6B | 17 | 0,008 | 26,7 | 0,005 | 37,0 |
| XRN1 | 2 | 0,011 | 24,3 | 0,006 | 38,7 |
| YTHDF1 | 5 | 0,017 | 5,3 | 0,010 | 5,9 |
| YTHDF2 | 5 | 0,028 | 4,3 | 0,007 | 7,3 |
| YTHDF3 | 5 | 0,013 | 5,7 | 0,005 | 10,8 |
| ZC3H14 | 6 | 0,022 | 11,3 | 0,005 | 19,3 |
